# HERMES: a molecular formula-oriented method to target the metabolome

**DOI:** 10.1101/2021.03.08.434466

**Authors:** Roger Giné, Jordi Capellades, Josep M. Badia, Dennis Vughs, Michaela Schwaiger-Haber, Maria Vinaixa, Andrea M. Brunner, Gary J. Patti, Oscar Yanes

**Author notes:** Corresponding author: Oscar Yanes, PhD, Department of Electronic Engineering, Universitat Rovira i Virgili, Avinguda Països Catalans, 26, 43007 Tarragona, Spain.

## Abstract

Comprehensive metabolome analyses are hampered by low identification rates of metabolites due to suboptimal strategies in MS and MS2 acquisition, and data analysis. Here we present a molecular formula-oriented and peak detection-free method, HERMES, that improves sensitivity and selectivity for metabolite profiling in MS and structural annotation in MS2. An analysis of environmental water, *E. coli*, and human plasma extracts by HERMES showed increased biological specificity of MS2 scans, leading to improved mass spectral similarity scoring and identification rates when compared to iterative data-dependent acquisition (DDA). HERMES is available as an R package with a user-friendly graphical interface to allow data analysis and interactive tracking of compound annotations.

## INTRODUCTION

A single LC/MS-based metabolomic experiment generates millions of three-dimensional (*m/z*, retention time, intensity) data points that can be annotated and quantified into thousands of metabolite features. However, most features are either redundant ions caused by ionization-related phenomena such as cation/anion adduction, multimerization and in-source fragmentation, or unknown contaminants and artifacts^1,2^. Moreover, conventional untargeted metabolomic experiments lead to highly heterogeneous chromatographic peak shapes, which negatively affect the performance of peak detection^3^ and grouping/annotation algorithms in MS1 mode^4^. These characteristics of MS1 data, in turn, negatively impact MS2 acquisition methods used for metabolite identification. In data-dependent acquisition (DDA) mode, MS2 spectra are automatically collected for precursor ions that exceed a predefined intensity threshold. The selection of precursor ions is a stochastic event suffering from low analytical reproducibility and favouring the selection of the most abundant, but not necessarily biologically relevant, ions. In data-independent acquisition (DIA) methods, multiple precursor ions, including redundant and biologically irrelevant ions, are simultaneously fragmented, often generating a series of complex convoluted MS2 spectra. Despite the emergence of new software to reconstruct the link between precursors and their fragments through mass spectral deconvolution^5,6^, MS2 spectral quality and matching scores to reference spectra are generally poorer in DIA compared to DDA^7^.

## RESULTS

Here we present HERMES, a novel experimental method and computational tool that improves the selectivity and sensitivity for comprehensive metabolite profiling in MS1, and identification in MS2. HERMES replaces the conventional untargeted metabolomic workflow that detects and annotates peaks^8,9^, for an inverse approach that directly interrogates raw LC/MS1 data points (i.e., scans) by using a comprehensive list of unique molecular formulas selected by the user. These are retrieved from large compound-centric databases (e.g., HMDB, ChEBI, NORMAN)^10–12^, genome-scale metabolic models, or specific metabolic pathways. Each molecular formula generates multiple ‘ionic formulas’ by adding or subtracting atoms from common adduct ions (Fig. 1). The resulting ionic formulas (on the order of 10^4^-10^5^ from a database such as HMDB) are searched against millions of data points in an LC/MS1 experiment. HERMES calculates the theoretical isotopic pattern of each ionic formula based on a predefined experimental mass resolution value (Suppl. Fig. 1). The number of collisions between monoisotopic ionic formulas vary according to the experimental mass error (i.e., the smaller the error, the larger the percentage of non-overlapping ionic formulas; Suppl. Fig. 2). An LC/MS1 data point contains *m/z* and intensity information in a wide mass range (e.g., *m/z* 80 to 1,000) for a given instant of time. HERMES solves the limitations of peak detection by finding a series of scans, named SOI (Scans Of Interest), which are defined as clusters of data points that match an ionic formula and are concentrated within a short period of time (see Methods). SOI shapes do not necessarily fit a Gaussian-like function, as assumed in basic chromatography theory, making the process independent of the heterogeneous peak shapes commonly observed in LC/MS1 experiments from complex mixtures. SOIs are then filtered in three steps: (i) blank subtraction from the sample based on a convolutional neural network (Suppl. Fig. 3a), (ii) adduct and isotopologue grouping according to the similarity of their elution profiles (Suppl. Fig. 3b), and (iii) in-source fragment (ISF) annotation by using publicly available low-energy MS2 data (Suppl. Fig. 3c) extending on Domingo-Almenara et al.^13^. Finally, users can prioritize the SOIs that will constitute the inclusion list (IL) for targeted MS2 acquisition based on the following criteria: type and number of adducts, minimum intensity, isotopic fidelity, and a maximum number of overlapped precursors at any time range, which together determine the total number of MS2 runs. According to the MS2 acquisition settings, each entry in the IL may be associated with one or multiple MS2 scans: if there are more than five continuous scans, HERMES provides an optional deconvolution step (adapted from CliqueMS^14^) that resolves co-eluting compounds (Suppl. Fig. 4); if there are fewer scans, HERMES selects the most intense scan. The resulting curated MS2 spectra can either be identified within HERMES or exported as .mzML, .msp, or .mgf files to be used in other identification software such as NIST MS Search, SIRIUS^15,16^, or GNPS^17^.

**Figure 1.**
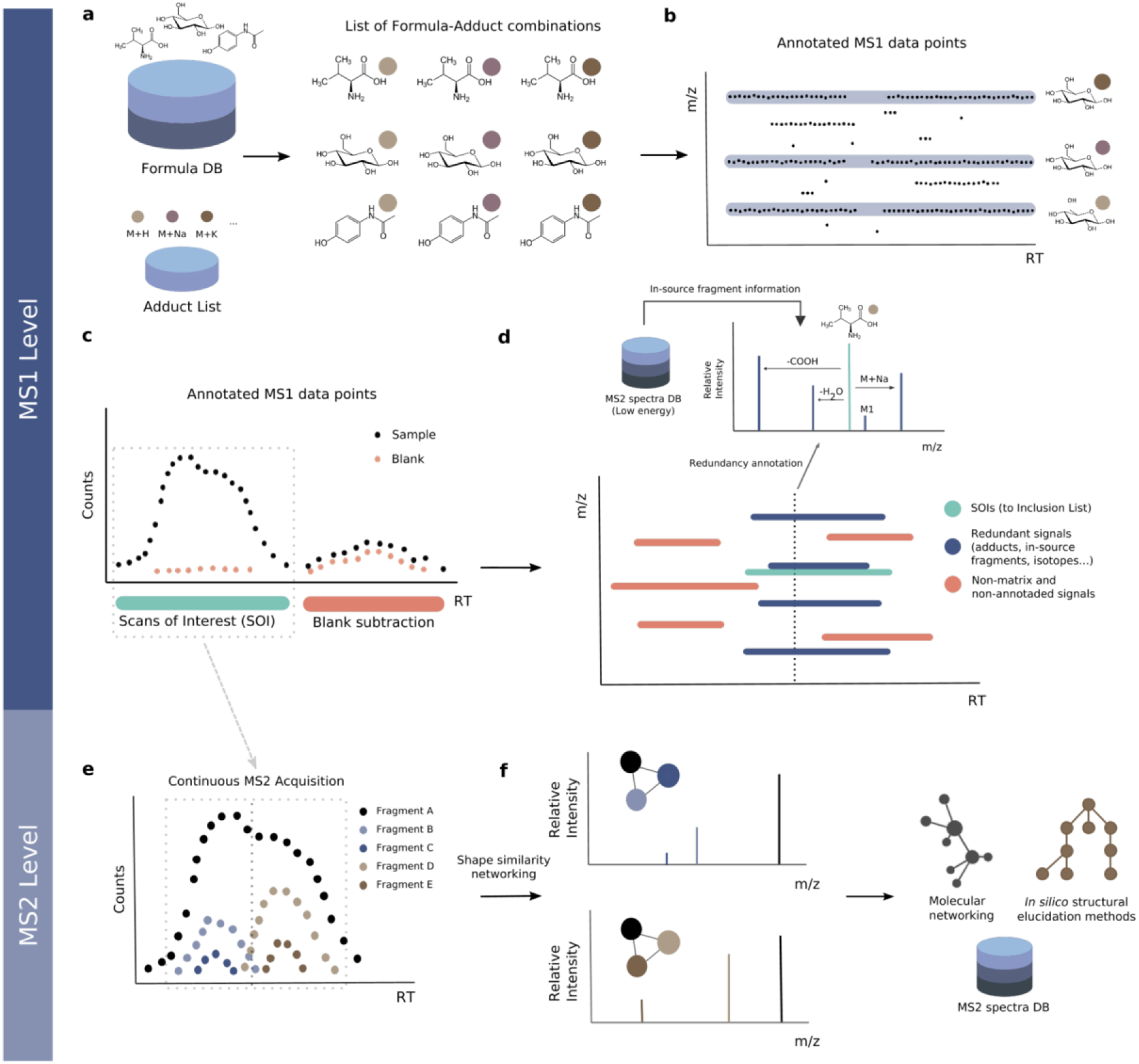
The HERMES workflow. (a) A context-specific database of molecular formulas and MS adducts generates a list of ionic formulas. (b) LC/MS1 data points are interrogated against all m/z ions corresponding to the ionic formulas and their isotopes. (c) Points with the same m/z annotation are grouped by density into retention time (RT) intervals called Scans of Interest (SOI). SOIs with similar shape and intensity in a blank sample are removed. (d) SOIs corresponding to different adducts of the same formula are grouped by their chromatographic elution profile. Similarly, in-source fragments are annotated based on low intensity MS2 spectra of molecules with the same formula. The result is an inclusion list (IL) of sample-specific and non-redundant precursor ions that will be monitored in a posterior MS2 experiment. (e) The IL entries are acquired continuously along the defined RT interval and HERMES groups the resulting fragment elution profiles. (f) This results in deconvoluted spectra that can be queried against an MS2 database or exported to be used in alternative identification workflows.

HERMES is available as an R package (RHermes) and comes with an R Graphical User Interface (GUI) to allow data analysis, tracking of compound annotations, and visualization (Suppl. Fig. 5). RHermes accepts both CSV and XLS/XLSX files as valid molecular formula lists and can extract formulas from selected KEGG pathways for a given organism. The running time, including blank subtraction and IL generation, is <10 minutes on a six-core, 2.9 GHz CPU.

HERMES has been validated by using three (bio)chemically relevant samples of increasing complexity: (i) water collected from a canal in Nieuwegein (Netherlands), (ii) *E*.*coli*, and (iii) human plasma extracts. The canal water was spiked with 86 common environmental contaminants at 1 µg/L (Suppl. Table 1) and analyzed by RP/LC (C18) coupled to an Orbitrap in positive (pos) and negative (neg) ionization mode operating at 120,000 resolution. Using 118,820 (pos) and 46,809 (neg) ionic formulas calculated from 24,696 unique molecular formulas in the NORMAN database, HERMES detected and annotated all spiked compounds at the MS1 level. Certain ionic formula collisions, particularly those involving Cl, Br, S, or K, were automatically resolved by matching experimental isotopic patterns to the expected ones. This is the case, for example, of the [M+H]^+^ ion of chloridazon and the [M+K]^+^ ion of 2-amino-alpha-carboline, which overlapped at 0.27 ppm (Suppl. Fig. 6). In-source fragments that could be wrongly associated with ionic formulas were also annotated by using low-energy MS2 spectra when available. The output was a curated IL of 474 (pos) and 129 (neg) selective entries for targeted MS2 (Suppl. Fig. 7).

Next, a reference *E. coli* cell extract (Cambridge Isotope Laboratories) was analysed by HILIC coupled to an Orbitrap in positive and negative ionization mode. LC/MS1 data were analysed by HERMES by using 12,010 (pos) and 4,876 (neg) ionic formulas calculated from 2,463 unique molecular formulas obtained from the *Escherichia coli* Metabolome Database (ECMDB) and KEGG database. Interestingly, HERMES annotated ionic formulas for 25% (pos) and 22% (neg) of all data points acquired by the mass spectrometer (Fig. 2a and Suppl. Fig. 8a). In comparison with XCMS, a commonly used open-source LC/MS1 processing data tool in untargeted metabolomics^9,18^, 4.5% of all acquired data points were associated with an XCMS peak, 1.5% of data points in XCMS peaks matched an ionic formula from ECMDB and KEGG database, and only 0.7% of data points in XCMS peaks were represented in the final SOI list after blank subtraction, isotopic fidelity, and ISF removal. The outcome of HERMES was 2,058 (pos) and 1,081 (neg) SOIs that led to a curated IL of 1,251 and 661 entries for targeted MS2, respectively.

**Figure 2.**
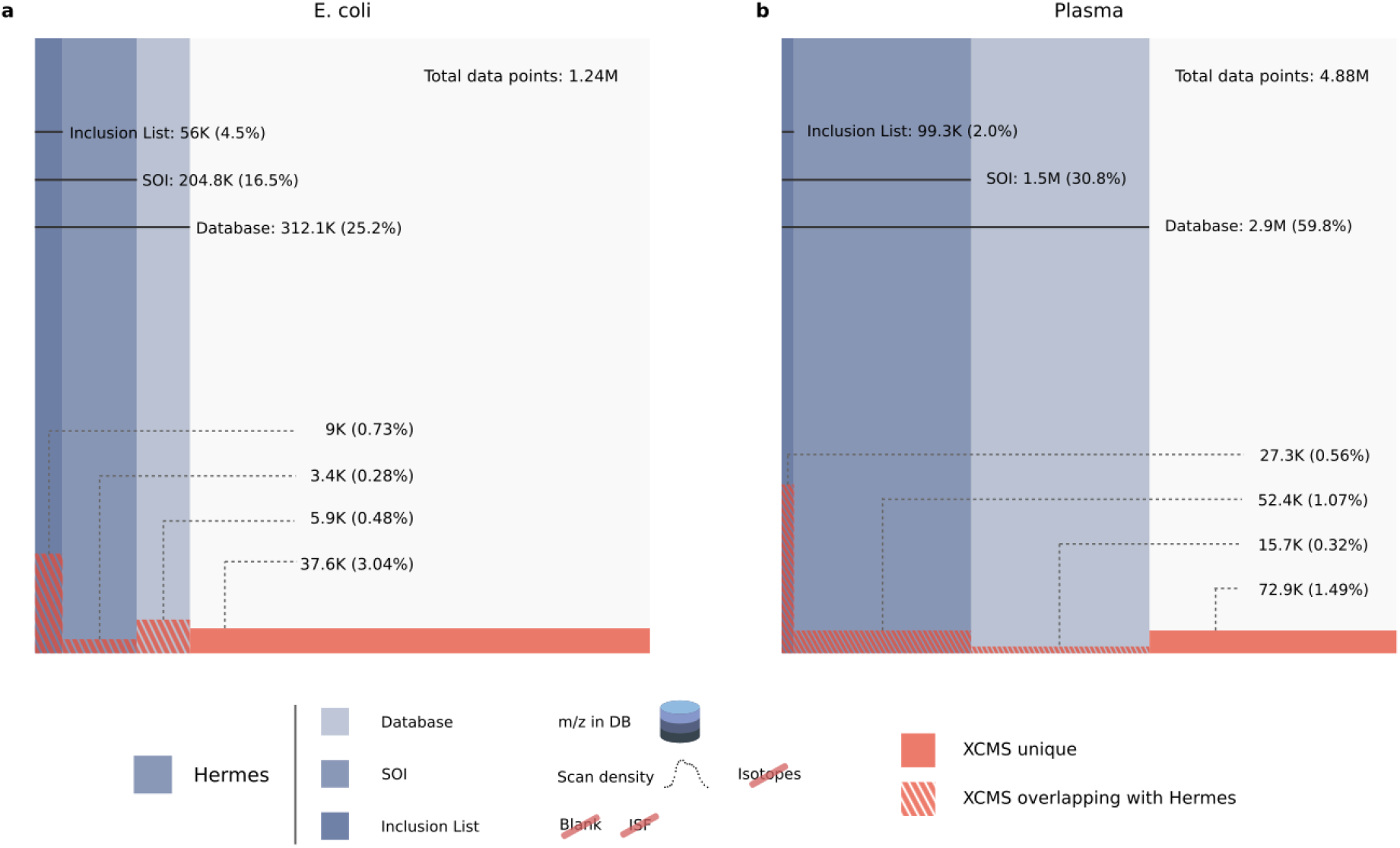
Venn-like diagram of the distribution of LC/MS1 data points in different steps of the HERMES workflow and XCMS peak-associated points. a) *E. coli* extract. b) Plasma extract. Database: Refers to all data points whose m/z matches with any m/z calculated from the ionic formula database (including isotopes). SOI: monoisotopic (M0)-annotated data points that are in Database and are also present in a SOI list that does not include blank subtraction nor any filtering. Inclusion List: data points present in Database and SOI kept through the blank subtraction, isotopic filter and ISF removal steps. Percentages refer to the total number of LC/MS1 data points. Positive ionization mode. On average, ∼78% of data points in the inclusion list could not be annotated as a peak by XCMS. Conversely, ∼84% scans annotated as a peak by XCMS could either not be matched to an ionic formula, were not specific of the sample or were associated with redundant signals.

The *E. coli* extract was also analysed by iterative DDA under identical analytical conditions. Remarkably, 68% of DDA scans could not be annotated as the monoisotopic signal by any ionic formula from ECMDB and KEGG database (Fig. 3a), which indicates their exogenous or artefactual origin. After filtering out DDA precursor ions that were classified as SOIs in the blank sample, redundant adducts, and ISF by HERMES; only 16% of the DDA scans matched with any monoisotopic ionic formula in the inclusion list. In addition, HERMES included 571 inclusion list entries (46.5% of the total) that were not triggered by DDA.

**Figure 3.**
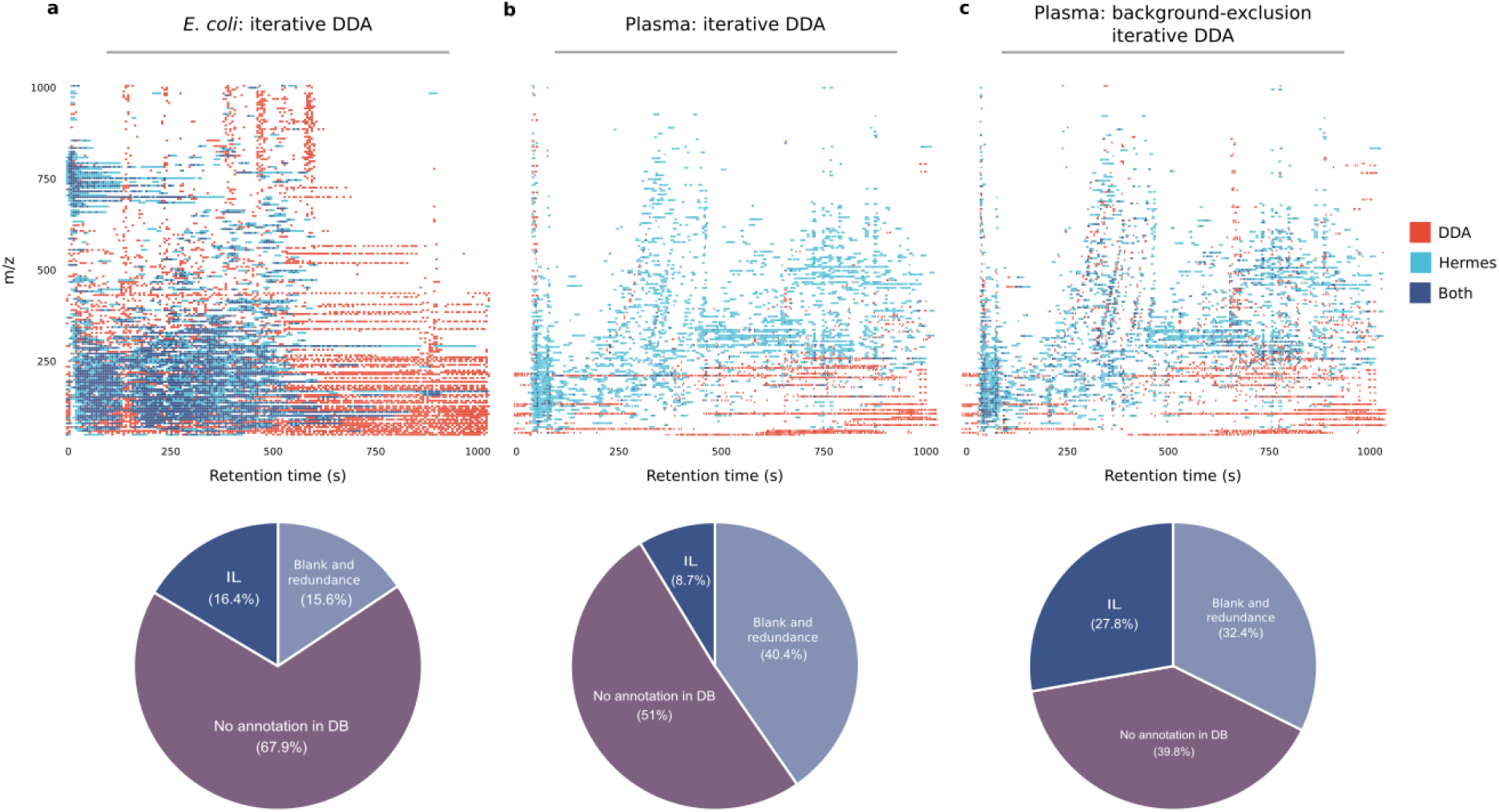
Distribution of MS2 scans acquired by HERMES and iterative DDA. a) Unlabeled *E. coli* and b) human plasma samples acquired by iterative DDA. c) Human plasma sample acquired by iterative DDA with background-exclusion. The acquired scans have been binned into 5Da-5s intervals. The precursor m/z of DDA scans have been queried into the corresponding ionic formula m/z database with a 3 ppm mass error tolerance. Scans annotated in the database were further classified according to whether the m/z and retention time of the scans could be matched to the HERMES inclusion list or not. Percentages in the pie-charts refer to the total number of acquired DDA MS2 scans.

To confirm the biogenic specificity of the MS2 scans in HERMES, a reference ^13^C-labeled (at ≥98% from uniformly ^13^C-labeled glucose) *E*.*coli* credentialing extract was analysed under identical LC/MS1 conditions. For each selected precursor ion in the unlabeled *E*.*coli* sample, we calculated its fractional contribution (FrC)^19–21^ and the monoisotopic ratio score (MIRS) by using the analogue ^13^C-labeled sample (see Methods). A metabolite with *n* carbon atoms can have zero (FrC=0) to *n* (FrC=1) of its carbon atoms labeled with ^13^C. In turn, similar intensity of the monoisotopic ion in the unlabeled and ^13^C-labeled *E*.*coli* extracts indicates no isotopic enrichment (MIRS=0), whereas loss of intensity in the ^13^C-labeled sample is associated with enrichment (MIRS=1). Around 63% of inclusion list entries in HERMES were associated with highly ^13^C-enriched metabolites (FrC and MIRS>0.5), proving the biosynthetic origin of these ions (Fig. 4a-c). These are mainly associated with abundant ions, while unlabeled precursors relate more frequently to low-abundant ions (Suppl. Fig. 9a,b). In contrast, only 20% of all DDA scans were associated with ^13^C-labeled and annotated precursors from ECMDB and the KEGG database, pointing to ions also present in the blank sample as the main source of unlabeled precursors (Fig. 4d-f). ^13^C-labeled precursors in DDA corresponded to highly abundant ions that were also covered by IL entries in HERMES (Suppl. Fig. 9c).

**Figure 4.**
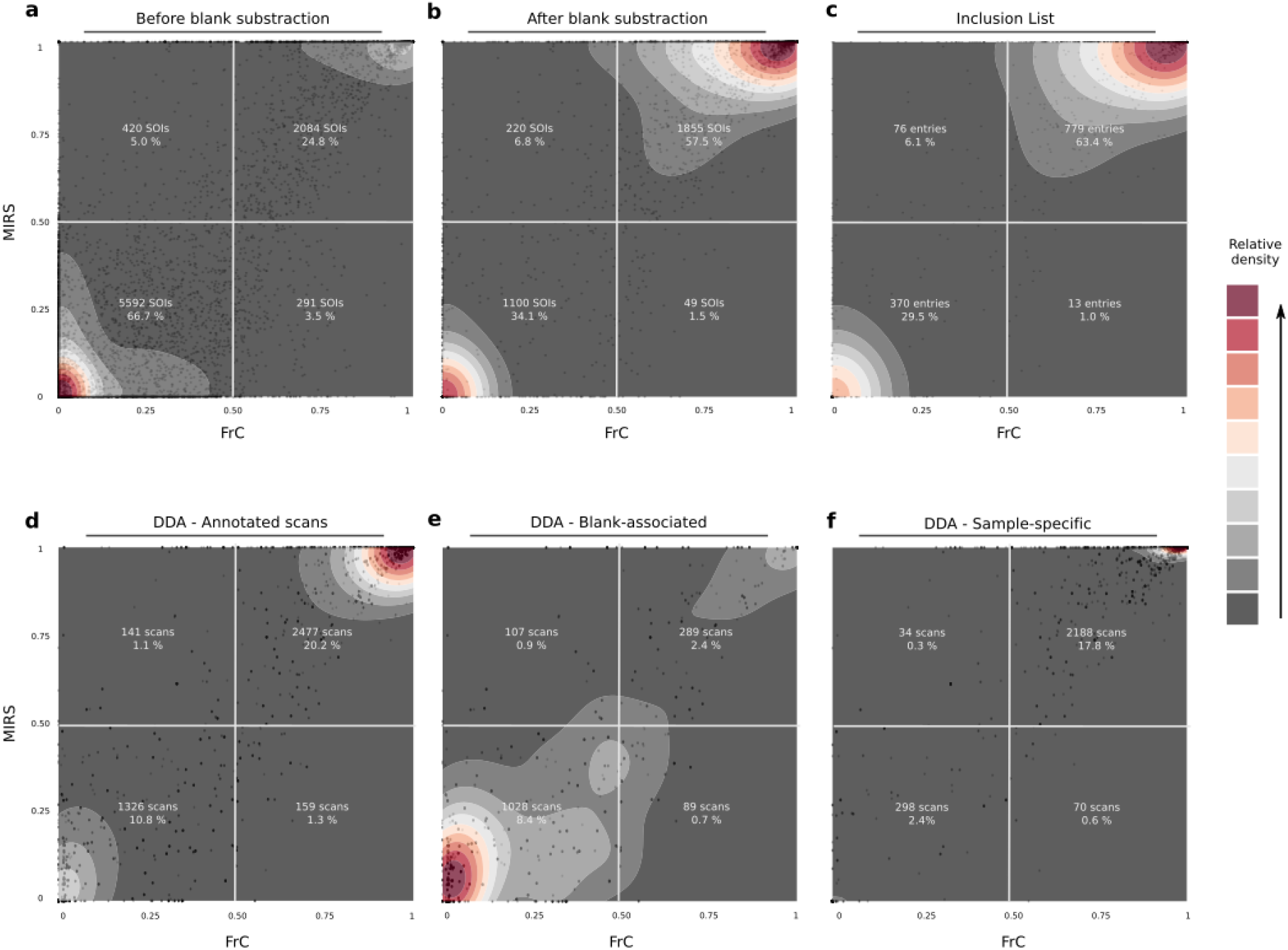
^13^C-enrichment analysis in the labeled *E*.*coli* sample. Each panel represents a scatterplot of two independent isotopic enrichment scores —FrC (Fractional Contribution) and MIRS (MonoIsotopic Ratio Score)─; and an overlaid density estimation. a) Distribution of SOIs before applying the blank subtraction filtering in HERMES. b) Same SOI list after removing most blank-related SOIs. c) SOIs in the MS2 inclusion list after removing redundant signals from b). d) Iterative DDA scans that could be matched to any m/z of the ionic formula database. e) DDA scans associated with SOIs removed during the blank subtraction step from a) to b). f) DDA scans associated with SOIs conserved during the blank subtraction. Percentages in a), b) and c) correspond to the total number of SOIs and inclusion list entries, accordingly, while percentages in d), e) and f) correspond to the total number of acquired DDA scans.

The biogenic specificity of HERMES resulted in higher similarity scores by mass spectral matching in databases (MassBankEU, MoNA, HMBD, Riken, NIST14, mzCloud)^22^ than iterative DDA (see Methods). HERMES provided nearly double the number of confident structural metabolite annotations than iterative DDA (Fig. 5a and Suppl. Fig. 10a). The higher identification rate of HERMES was validated by using alternative spectral similarity and distance metrics (Suppl. Fig. 11). A fraction of the ^13^C-labeled compounds, however, could not be identified due to low intensity SOIs and/or the lack of reference spectra in databases. For the former, setting the maximum ion injection time at high values (1,500 ms) improved sensitivity and MS2 spectral quality in HERMES, resulting in more informative fragments and better spectral matching (Suppl. Fig. 12). Furthermore, we identified unlabeled metabolites (FrC=0) in the ^13^C-labeled *E*.*coli* sample, such as choline, that we attribute to contaminants of the minimal growth medium that could not properly be removed by blank subtraction.

**Figure 5.**
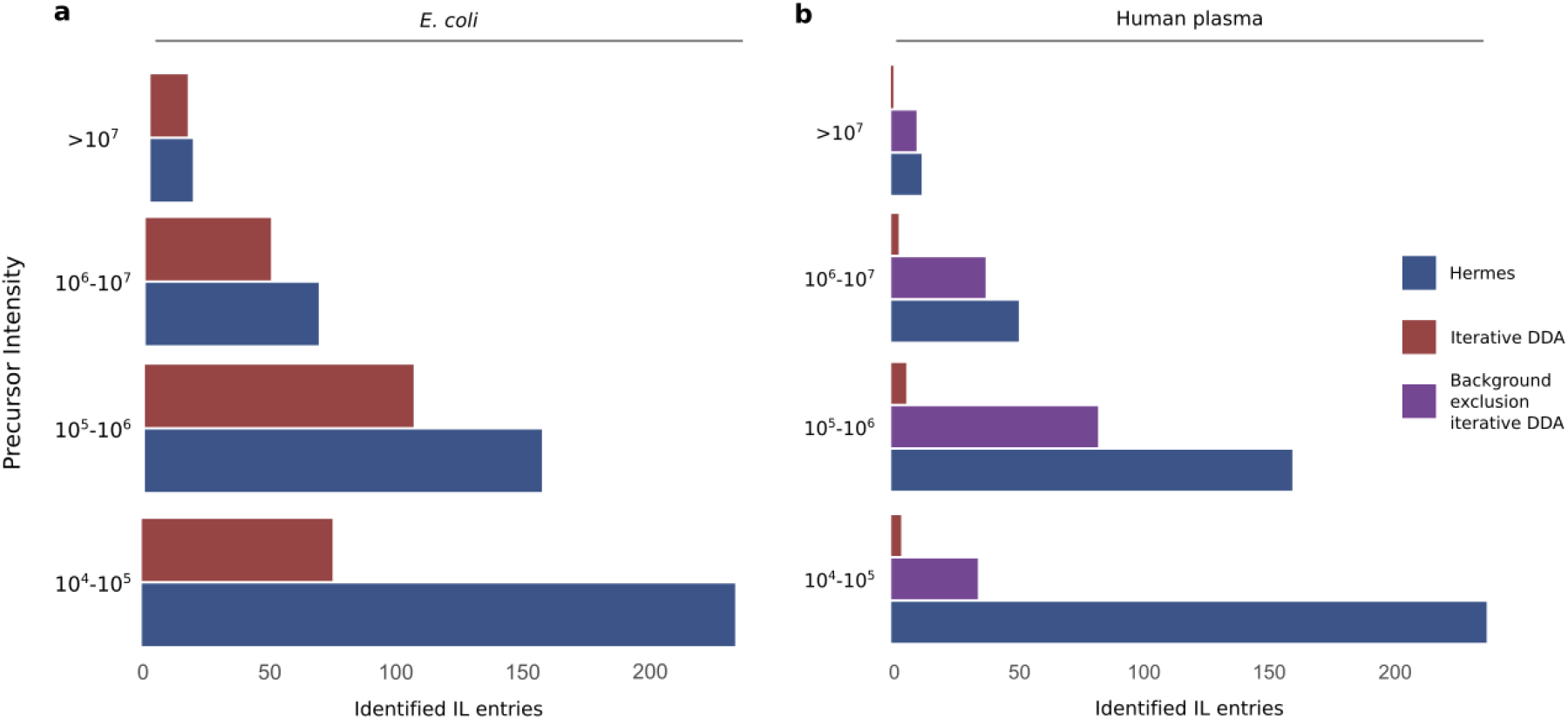
Identified inclusion list entries according to the MS1 precursor intensity. An inclusion list (IL) entry is considered identified if at least one MS2 scan associated with it has a compound hit in the reference MS2 database with either cosine score > 0.8 (in-house database from MassBankEU, MoNA, Riken and NIST14 spectra), or Match > 90 and Confidence > 30 (mzCloud). Positive ionization data. a) *E. coli* extract. b) Human plasma extract.

Finally, we used a human plasma extract to compare HERMES and iterative DDA, with and without background exclusion^23^. Here we used 23,797 unique molecular formulas from the HMDB and Chemical Entities of Biological Interest (ChEBI) database to explore virtually all known exogenous and endogenous small molecules in this biofluid. HERMES generated 110,387 and 46,973 ionic formulas that covered 60% and 14% of all data points acquired by LC (RPC18)-Orbitrap MS in positive and negative ionization mode, respectively (Fig. 2b and Suppl. Fig 8b). Consistent with the pattern observed for *E. coli*, 3.8% of all acquired data points were associated with an XCMS peak. Only 0.5% of the data points in these peaks matched with an ionic formula from HMDB or ChEBI and were present in the inclusion list. Again, more than half of DDA precursors could not be annotated as monoisotopic ionic formulas from HMDB and ChEBI without blank subtraction (Fig. 3b). As expected, the overlap between HERMES and DDA increased upon background exclusion in iterative DDA, increasing to 29% of the number of common MS2 scans (Fig. 3c). Yet, the number of confident structural metabolite identifications with HERMES was more than three times greater than DDA because of the larger coverage of sample-specific and low abundant precursor ions (Fig. 5b and Suppl. Fig. 10b).

## DISCUSSION

Our results demonstrate that a conventional LC/MS-based untargeted metabolomic experiment can contain up to ∼50 times more non-specific and redundant data points than sample-specific and selective ones, which can account for as much as 90% of the MS2 acquisition run time in an iterative DDA experiment. Current untargeted metabolomic approaches are unable to properly annotate the large number of ‘junk’ MS and MS2 signals, leading to false-positive identifications and an overall low number of identified metabolites. HERMES solves this problem by implementing a broad scope and molecular formula-oriented method that improves MS2 coverage by optimizing MS2 acquisition time focusing on sample-specific, MS1 pre-annotated, and biologically relevant compounds, thereby increasing the quality of MS2 spectra and the number of identified metabolites. The use of molecular formulas restricts the range of known and unknown chemical structures for *in silico* MS2 fragmentation tools, avoiding the loss of possible unknown isomeric forms in a sample, and facilitating *de novo* MS2 annotation. HERMES, in addition, provides maximum experimental flexibility by allowing users to add new molecular formulas not reported in public databases, including *in silico* secondary metabolism prediction^24–26^ such as environmental microbial degradation, biotransformations of gut and soil/aquatic microbiota, or small peptides such as dipeptides and tripeptides. Finally, future developments should provide optimized maximum ion injection time and collision energies for each IL entry to reduce the number of MS2 scans required and improve the quality of MS2 spectra for all inclusion list entries, particularly for low intensity SOIs, as current Orbitrap mass spectrometers only allow fixed injection times.

## METHODS

### Materials

LC/MS-grade acetonitrile, water, isopropanol, and methanol (Burdick & Jackson) were purchased from Honeywell (Muskegon, MI). LC/MS-grade ammonium acetate and ammonium hydroxide were purchased from Sigma-Aldrich (St. Louis, MO). TraceSELECT Fluka brand ammonium phosphate (monobasic) was purchased from Honeywell (Muskegon, MI). Dried down metabolic extracts of *E. coli* were purchased from Cambridge Isotope Laboratories (MSK-CRED-DD-KIT). Spike-in compounds (Suppl. Table 1) were purchased from Sigma-Aldrich (Zwijndrecht, The Netherlands), LGC Standards (Wesen, Germany) and Toronto Research Chemicals (Toronto, ON).

### Sample preparation

#### Environmental water

Surface water was obtained from the Lekkanaal at Nieuwegein (The Netherlands). The spike-in compounds were added to the surface water sample to a final concentration of 1 µg/L. Subsequently, the sample was filtered using Phenex™ reversed cellulose 15 mm Syringe Filters 0.2u (Phenomenex, Torrance, USA) and transferred to a LC autosampler vial.

#### E. coli

Dried down *E. coli extracts (*unlabeled and uniformly ^13^C-labeled) were reconstituted in 100 μL of acetonitrile:water (2:1), followed by 30 s vortexing, 5 min of sonication, and 30 s of vortexing.

#### Human plasma

Plasma aliquots (50 μL) were thawed at 4°C and briefly vortex-mixed. Proteins were precipitated by the addition of 200 μL cold acetonitrile/methanol/water (5:4:1, vol/vol) followed by 10 seconds vortex-mixing. Samples were subsequently maintained on ice for 30 min. After centrifugation (10 min, 15.200 rpm at 4°C), 100 μL of supernatant were transferred to a LC autosampler vial.

### LC-MS analysis

#### Environmental water and human plasma

Ultra-high performance LC (UHPLC)/MS was performed with a Thermo Scientific Vanquish UHPLC system interfaced with a Thermo Scientific Orbitrap Fusion Tribrid mass spectrometer operated in positive or negative ion mode. Reverse phase C18 liquid chromatography (RPLC) analysis was performed by using a Xbridge BEH C18 column (Waters, Etten-Leur, The Netherlands) with the following specifications: 150 mm x 2.1 mm, 2.5 μm. Mobile-phase solvents were composed of A = ultrapure water with 0.05% formic acid (v/v) and B = acetonitrile with 0.05% formic acid (v/v). The column compartment was maintained at 25 °C for all experiments. The following linear gradient was applied at a flow rate of 250 μL/min: 0-1 min: 5% B, 1-25 min: 5-100% B, 25-29 min: 100% B, 29.0-29.5 min 5% B followed by 4.5 min of re-equilibration phase. One μL of the human plasma extract was diluted in 100 μL of ultrapure water, and the injection volume was 100 μL for all experiments. Data were collected with the following settings: spray voltage, 3.0 kV and -2.5 kV in positive and negative mode, respectively; sheath gas, 40; auxiliary gas, 10; sweep gas, 5; ion transfer tube temperature, 300 °C; vaporizer temperature, 300 °C; mass range, 80-1000 Da; RF lens, 50%; resolution, 120,000 (MS1), 15,000 (MS/MS); AGC target, 2e5 (MS1), 5e4 (MS2); maximum injection time, 100 ms (MS1), 50 ms (HERMES), 50 ms (DDA); isolation window, 1.6 Da. The collision energy was 35% for HCD fragmentation. With every batch run, mass calibration was performed using Pierce ESI positive and negative ion calibration solution in order to obtain a mass error of <2 ppm.

#### E. coli

LC/MS was performed with a Thermo Scientific Vanquish Horizon UHPLC system interfaced with a Thermo Scientific Orbitrap ID-X Tribrid Mass Spectrometer (Waltham, MA). Hydrophilic interaction liquid chromatography (HILIC) analysis was performed by using a SeQuant ZIC-pHILIC column (Merck Millipore, Burlington, MA) with the following specifications: 150 mm x 2.1 mm, 5 μm. Mobile-phase solvents were composed of A = 20 mM ammonium bicarbonate, 0.1% ammonium hydroxide solution (25% ammonia in water) and 2.5 μM medronic acid in water:acetonitrile (95:5) and B = 95% acetonitrile, 5% water, 2.5 µM medronic acid. The column compartment was maintained at 40 °C for all experiments. The following linear gradient was applied at a flow rate of 250 μL min-1: 0-1 min: 90% B, 1-12 min: 90-35% B, 12.5-14.5 min: 25% B, 15 min: 90% B followed by 4 min of re-equilibration phase at 400 µL min-1 and 2 min at 250 µL min-1. The injection volume was 2 μL for all experiments. Data were collected with the following settings: spray voltage, 3.5 kV and -2.8 kV in positive and negative mode, respectively; sheath gas, 50; auxiliary gas, 10; sweep gas, 1; ion transfer tube temperature, 300 °C; vaporizer temperature, 200 °C; mass range, 70-1000 Da; RF lens, 60%; resolution, 120,000 (MS1), 15,000 (MS/MS); AGC target, 2e5 (MS1), 5e4 (MS2); maximum injection time, 200 ms (MS1), 35 ms (HERMES, unless otherwise stated), 100 ms (iterative DDA); isolation window, 1 Da. The collision energy was 35% for HCD fragmentation.

### Iterative DDA

#### E. coli

After the first DDA run, the raw data file containing MS/MS spectra was converted to an .MS2 file using MS Convert^27^ Next, the IEomics tool^28^ was used to generate the first exclusion list of features fragmented in the first DDA run. User inputs in the R script were RTWindow = 0.3 min, noiseCount = 25, MZWindow = 0.001. This procedure was repeated two times, which resulted in a total of three DDA data runs per polarity. The mass tolerance for exclusion lists was 5 ppm.

#### Plasma

An exclusion list of background ions was generated using the AcquireX workflow of Xcalibur data acquisition software (Thermo Fisher Scientific), by analyzing an ultrapure water sample. The exclusion list contains the exact mass, retention window and intensity (exclusion override factor = 3) of the excluded background ions. DDA was per performed for the top 6-8 most intense ions per full scan. Dynamic exclusion was used to prevent redundant acquisition of MS2 spectra for a selected precursor ion for 10 s, when two MS2 spectra were acquired within 20 s, resulting in a total of three DDA data runs per polarity. A mass tolerance of 5 ppm was used for the exclusion list and dynamic exclusion.

### HERMES algorithm

All analysis were performed using RHermes (version 0.99.0).

#### MS1 data processing

Theoretical isotopic patterns of each ionic formula were calculated by Envipat (version 2.4) and refined by RHermes, based on the predefined experimental mass resolution and mass accuracy values. Local resolution was calculated for each ionic formula as:

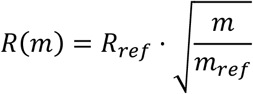

Using as input a set of mzML files, SOIs were detected by RHermes using two sets of 5s bins (offset by 2.5s) and required a minimum scan density of 30% of acquired scans.

Blank subtraction was performed using an heuristic prefilter (intensity ratio sample/blank > 3) and an artificial neural network trained with >3000 manually annotated sample/blank SOI pairs. Adduct and isotopologue grouping were performed using a cosine shape similarity score and required a cosine >0.8 and >0.85, respectively.

In-source fragment (ISF) annotation was performed using an in-house MS2 database consisting of MassBankEU, MoNA, HMBD, Riken and NIST14 spectra. Low intensity spectra (<20% HCD, <20eV CID) were selected according to each SOI formula annotation. Intense fragments (>20% of maximum intensity) m/z were then queried against the SOI list. Finally, the suspected ISF SOIs elution profiles were compared to the original SOI and a cosine similarity score was calculated.

#### MS2 data processing

The program exports the IL into a csv file used to generate the MS2 acquisition method. Acquired MS2 scans were linked to each IL entry; if >5 scans were acquired, a deconvolution algorithm was applied, where fragments m/z were grouped and split with a Centwave peak picking (peakwidth = c(5,60)). A cosine shape similarity score was applied to each pair of fragment peaks to generate a similarity network. Each network was then partitioned using a greedy algorithm from *igraph* (version 1.2.4.2) and resulted in a list of deconvoluted MS2 spectra. If fewer than 5 scans were acquired, the scan with the highest TIC was selected and the fragments were filtered by intensity (> 0.5% of maximum).

### XCMS data processing

LC-MS raw data files (ESI+ and ESI- modes) were converted to open standard format mzML using Proteowizard MS-convert^27^ and subsequently processed by HERMES and XCMS software^18^ (version 3.8.1). XCMS settings were: xcmsSet(method=“centWave”, ppm=8, peakwidth=c(1,60); Common data points between SOIs in HERMES and XCMS peaks were calculated by extracting the raw data points delimited by each XCMS peak (rt_min_< rt < rt_max_and mz_min_ < mz < mz_max_) and generating the set intersections using *dplyr* (version 1.0.4).

### Uniformly ^13^C-labeled *E. coli*

Fractional contribution (FrC) was calculated using the formula:

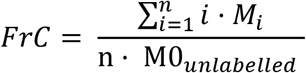

Where *M*_*i*_ is the intensity of the ^13^C_i_ isotope and n is the total number of carbon atoms in the molecule.

MonoIsotopic Ratio Score (MIRS) was calculated using the formula:

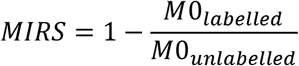

If MIRS is smaller than zero, it is set to zero so that all points range from 0 to 1.

### Identification by MS/MS

#### In-house DB

MS/MS spectra were obtained from MassBankEU, MoNA, HMBD, Riken and NIST14 databases. All fragment m/z were discretized into 0.01Da bins. Each spectrum precursor m/z was matched against the DB spectra m/z with a 0.01Da tolerance. For the HERMES matching, the reference spectra were further filtered according to the formula database used in the MS1 analysis. A cosine similarity score was calculated between the query and reference spectra and resulting hits were filtered by requiring a score > 0.8.

#### mzCloud DB

The processed HERMES MS2 spectra were exported to the mzML file format. The DDA files were directly imported through MassFrontier version 8.0 SR1 (Thermo Scientific) and matched against the mzCloud database using three component identification types: Identity, Similarity Forward and Similarity Reverse; with the following constraints: 4.0 Tolerance Factor and Match Ion Activation Type. The resulting hits were filtered by both Match and Confidence scores (requiring a score > 90 and > 30, respectively)

Identified IL entries (Figure 5 and Supp Figure 9) were calculated as number of IL entries that resulted in a valid hit (ie. high score) against either of the two databases. For DDA, this number was calculated by matching the precursor m/z and RT of the scans to the IL and then examining if (i) any of the scans have at least one valid hit against either of the two databases and (ii) any valid hit had a molecular formula present in the HERMES formula database.

All similarity metrics were calculated using the R package philentropy (version 0.4.0). MS2 spectra were discretized into 0.01Da bins and their fragment intensities scaled by the sum of the intensities, so that all calculated metrics were comparable across the spectra. The query spectra (both DDA and HERMES) were matched against the previously described In-house DB. For each query, all DB hits were grouped, taking the maximum similarity (cosine and fidelity) and the lowest distance (squared chord and topsoe). Additionally, HERMES hits were restricted to compounds with formulas present in the HERMES formula database. The corresponding plots were generated using ggplot2 (version 3.3.3).

## Data availability

Input mzML mass spectrometry data files and RMarkdown files are available at Zenodo with the accession number <u>4581662</u>.

## Code availability

The source code of RHermes is offered to the public as a freely accessible software package under the GNU GPL, version 3 license, and is available at https://github.com/RogerGinBer/RHermes.

## Acknowledgements

We gratefully acknowledge financial support by Ministerio de Educación y Formación Profesional (Spanish Government) to R.G. (2020-COLAB-00552). O.Y. was supported by Ministerio de Economía y Competitividad (MINECO) (BFU2014-57466-P), Spanish Biomedical Research Centre in Diabetes and Associated Metabolic Disorders (CIBERDEM), an initiative of Instituto de Investigación Carlos III (ISCIII), and the European Union’s Horizon 2020 program (MSCA-ITN-2015; 675610). We thank members of the Mil@b for helpful comments.

## Author contributions

RG and OY designed the research. RG, JC, JMB, MV and OY developed the computational method. DV and MSH performed LC-MS and MS/MS experiments. All authors applied and evaluated the method on biological samples. RG and OY wrote the manuscript, in cooperation with all authors.

## Competing interests

The authors declare no competing interests. A patent application for the method has been filled (P202030061).

## Supplemental Figures

**Supplementary Figure 1.**
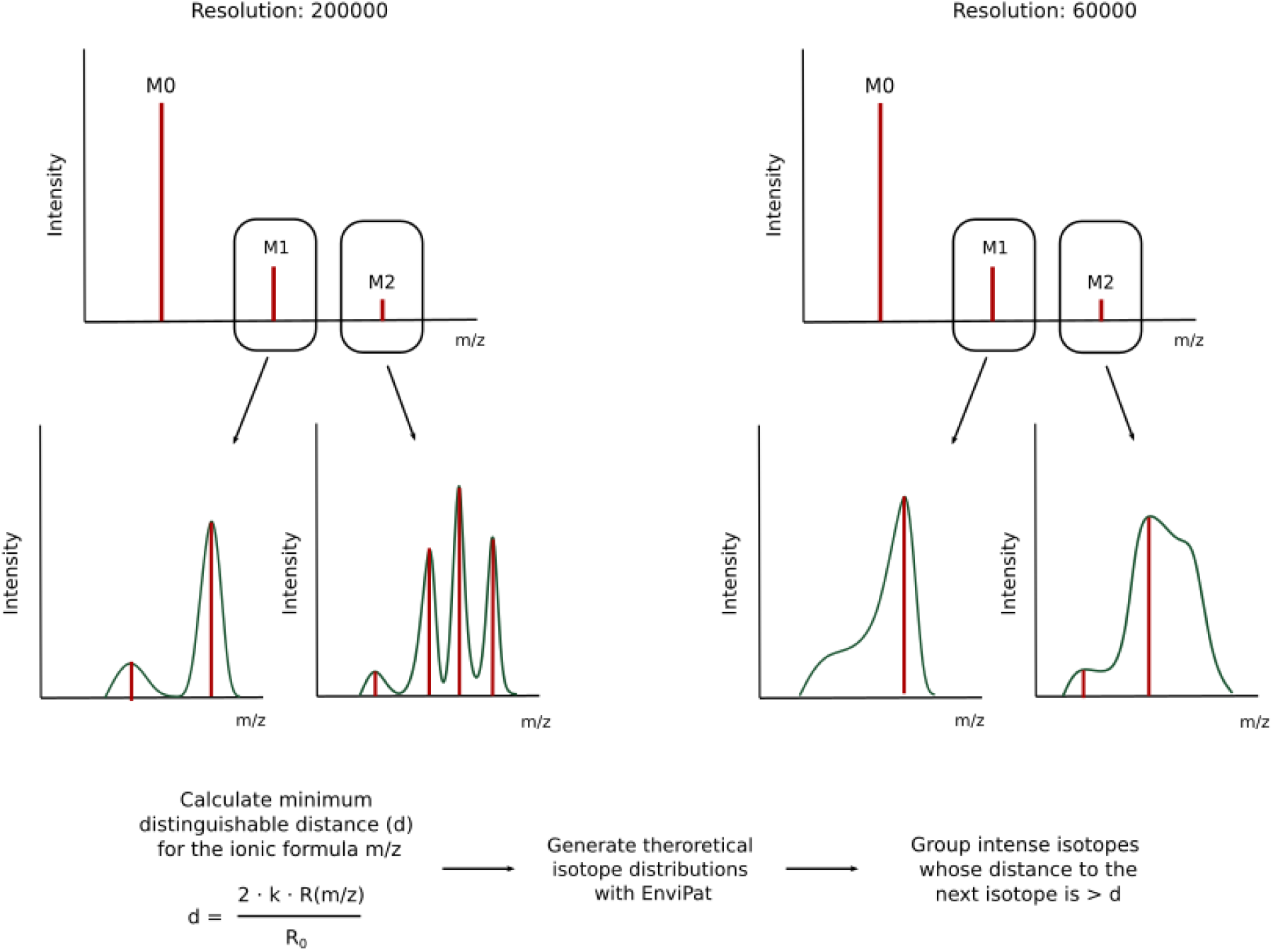
Calculation of the theoretical isotopic pattern of each ionic formula based on predefined experimental mass resolution values. By calculating a resolution-based parameter d, it is possible to estimate which close isotopologues are likely to be distinguishable in the acquired profile MS1 data and therefore present in the centroided data.

**Supplementary Figure 2:**
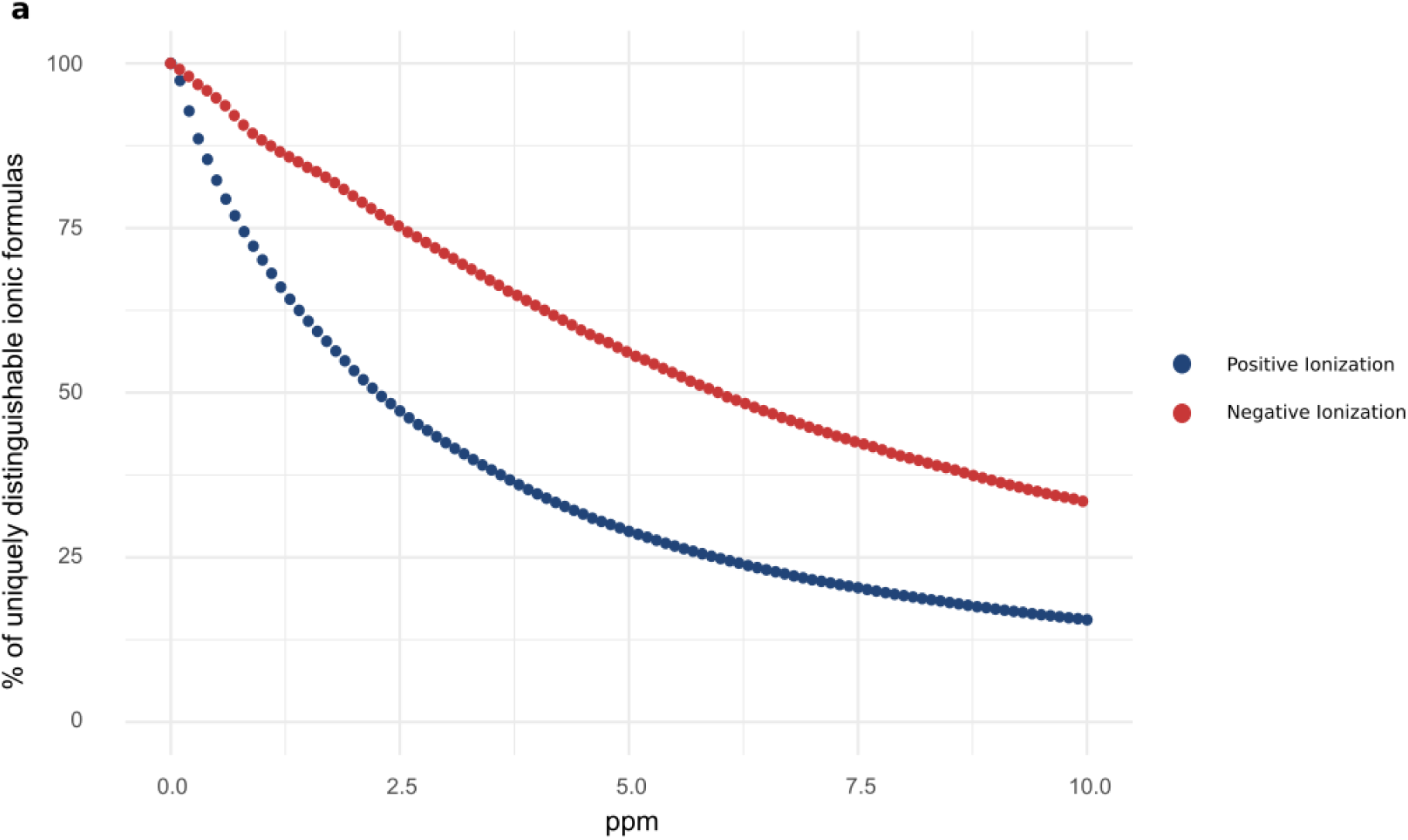
Ionic formula collisions from the NORMAN database (24,696 unique molecular formulas). Distribution of uniquely distinguishable ionic formulas. Blue: Positive ionization taking [M+H]^+^, [M+Na]^+^, [M+K]^+^, [M+NH4]^+^ and [M]^+^ adducts. Red: Negative ionization taking [M-H]^-^ and [M+Cl]^-^ adducts. As the ppm error of the instrument increases, the larger the percentage of overlapping ionic formulas.

**Supplementary Figure 3.**
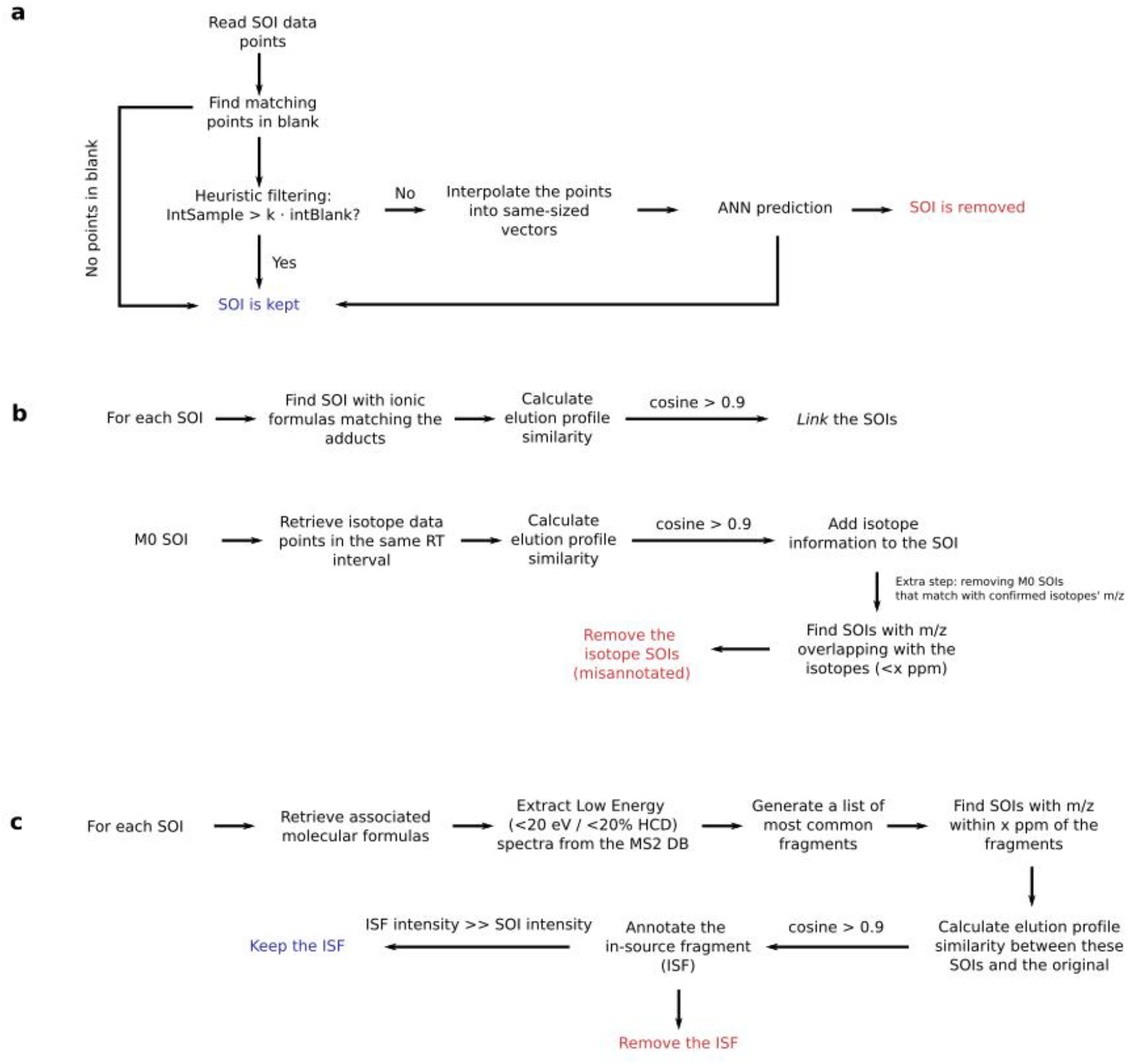
Schematic workflow of the different filtering steps in HERMES. a) Artificial neural network (ANN) for blank subtraction. b) Adduct and isotopologue grouping according to the similarity of their elution profiles. c) In-source fragment annotation, by using publicly available low-energy MS/MS data.

**Supplementary Figure 4.**
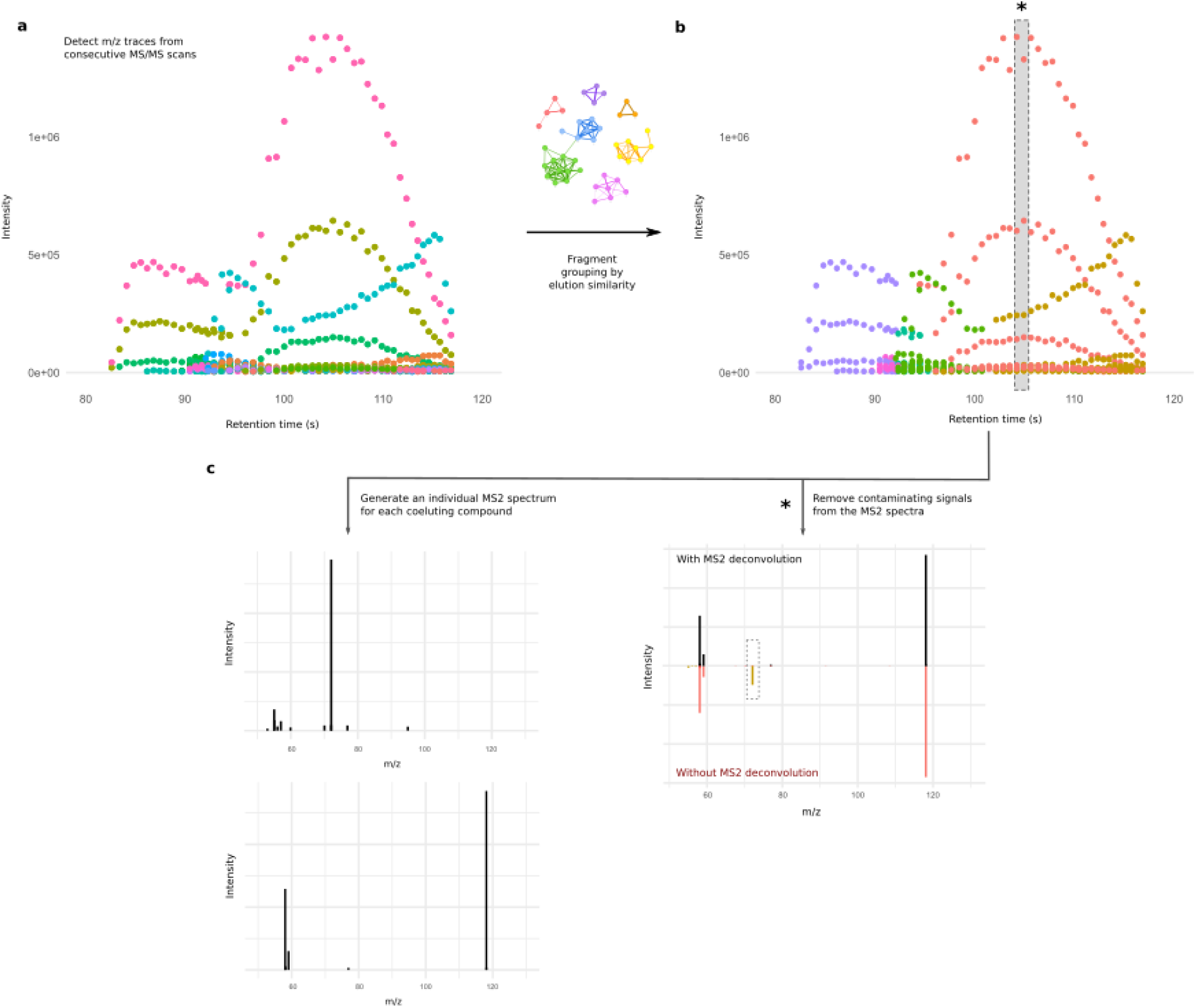
Continuous MS2 acquisition resolves co-eluting ionic species by comparing their fragment elution profile. a) All fragment ions from continuous MS2 scans are grouped according to their m/z. b) A loose peak-picking algorithm is applied and the resulting peaks are grouped according to their elution profiles, generating a similarity network that is split by a greedy clustering algorithm. c) This grouping yields a curated MS2 spectra for each coeluting species. (*) The shaded slice shows the impact of the algorithm on the resulting spectral quality. The delineated fragment in yellow has a different elution pattern than the rest and would contaminate the MS2 spectra if only one scan was acquired at the top of the peak. The grouping performed by HERMES confidently removes the contaminant ion and separates each group of fragments accordingly.

**Supplementary Figure 5.**
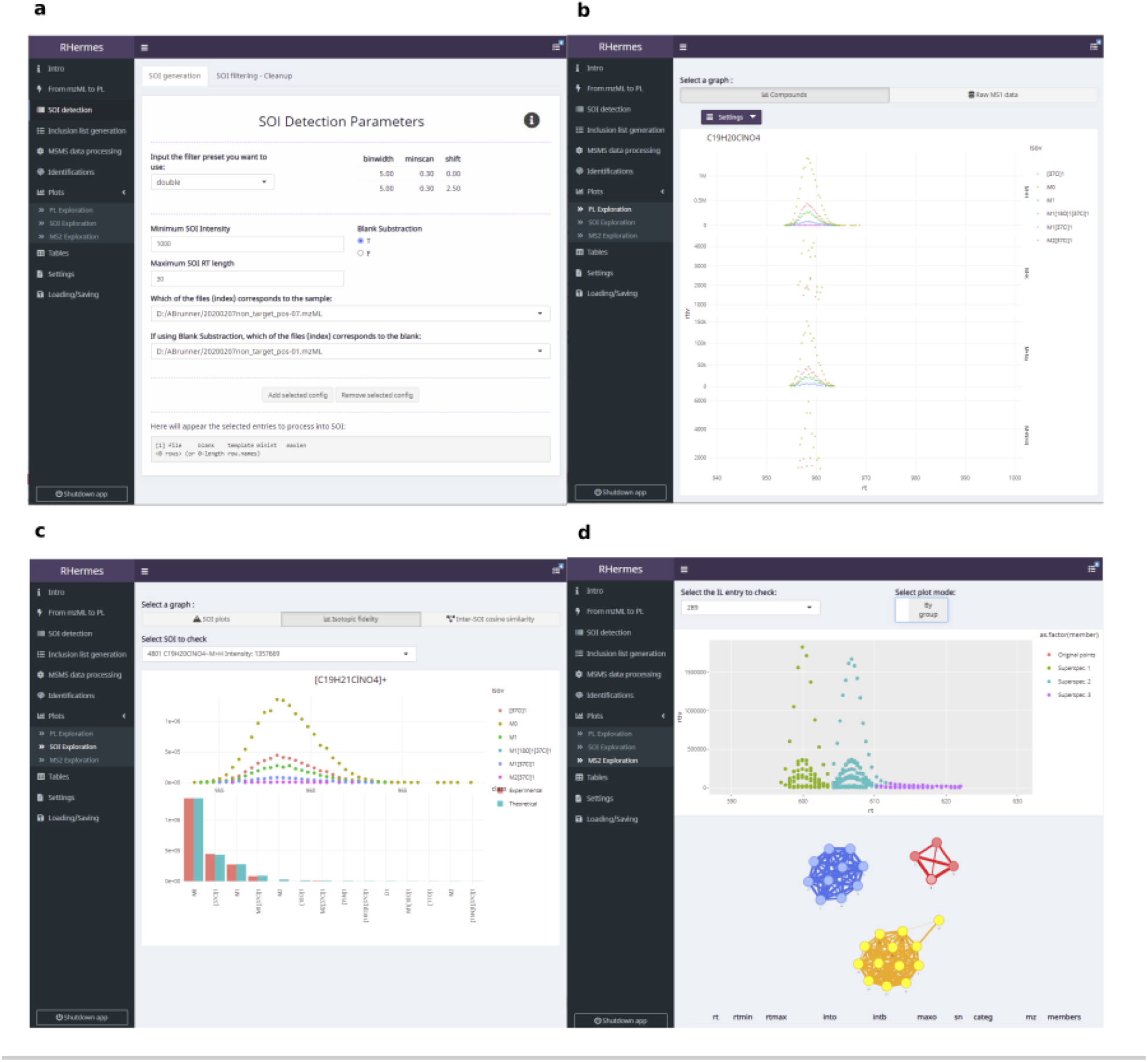
HERMES R Graphical User Interface (GUI). a) Point-and-click selection of SOI detection parameters, with detailed explanations on their usage and optimal values. b) Visualization of isotopic profiles of different adducts of the same formula. The formula can be inputted directly or inferred from the name of a compound chosen by the user. c) Isotopic fidelity exploration of selected SOIs. d) Visualization of the continuous MS2 deconvolution step. The user can check the fragment ion elution profiles from each inclusion list entry and how they are interconnected in the corresponding profile similarity network.

**Supplementary Figure 6.**
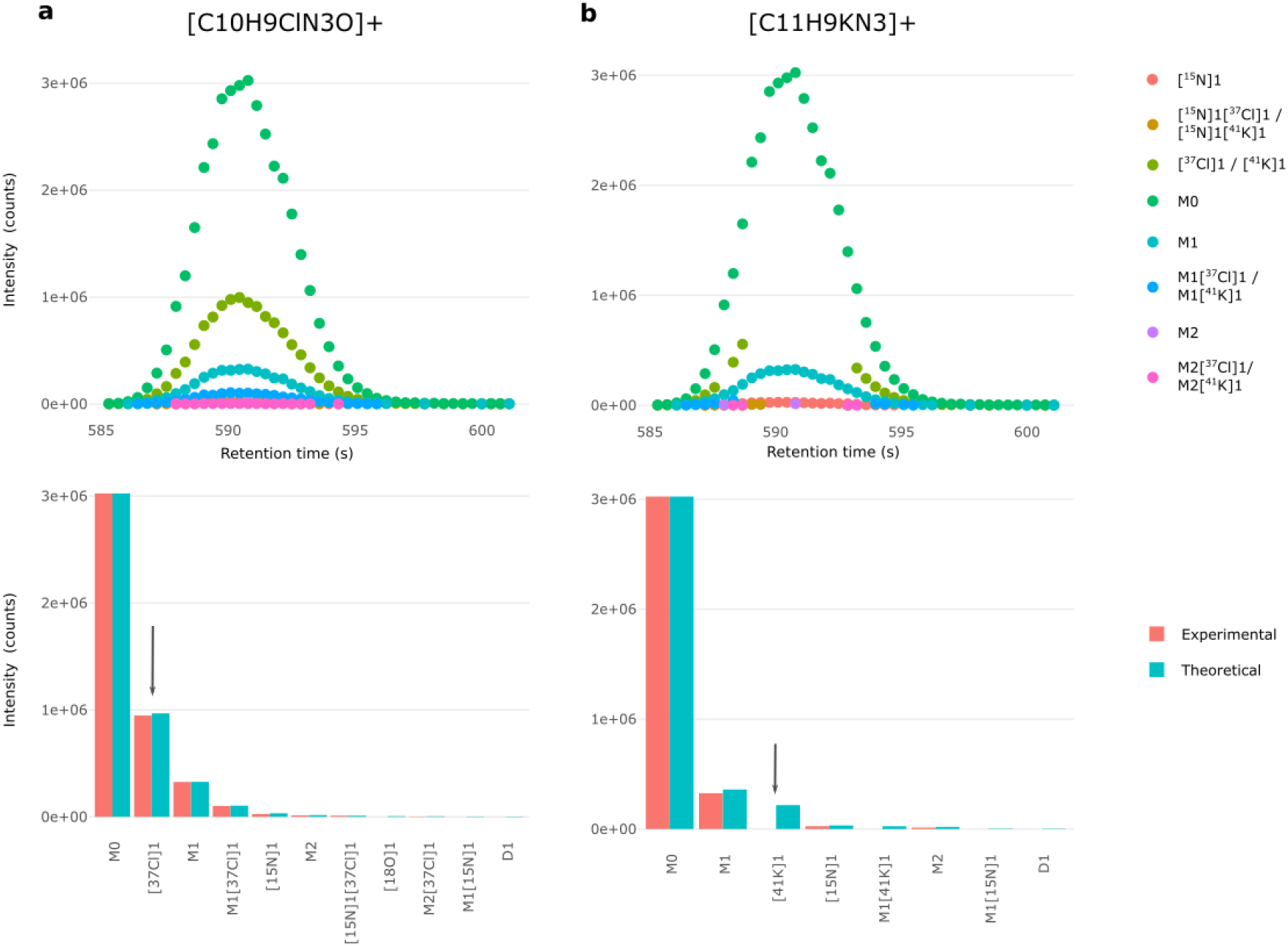
Discrimination of SOIs based on isotopic fidelity. a) [M+H]^+^ion of chloridazon and b) [M+K]^+^ ion of 2-Amino-alpha-carboline overlapping at 0.27 ppm. The arrows indicate the characteristic [^37^Cl] isotopologue present in chloridazon and the [41K] isotopologue absent in 2-Amino-alpha-carboline. The absence of characteristic isotopologue signals (Cl, Br, K, etc.) in intense SOIs results in a low isotopic fidelity score and the removal of such SOIs.

**Supplementary Figure 7.**
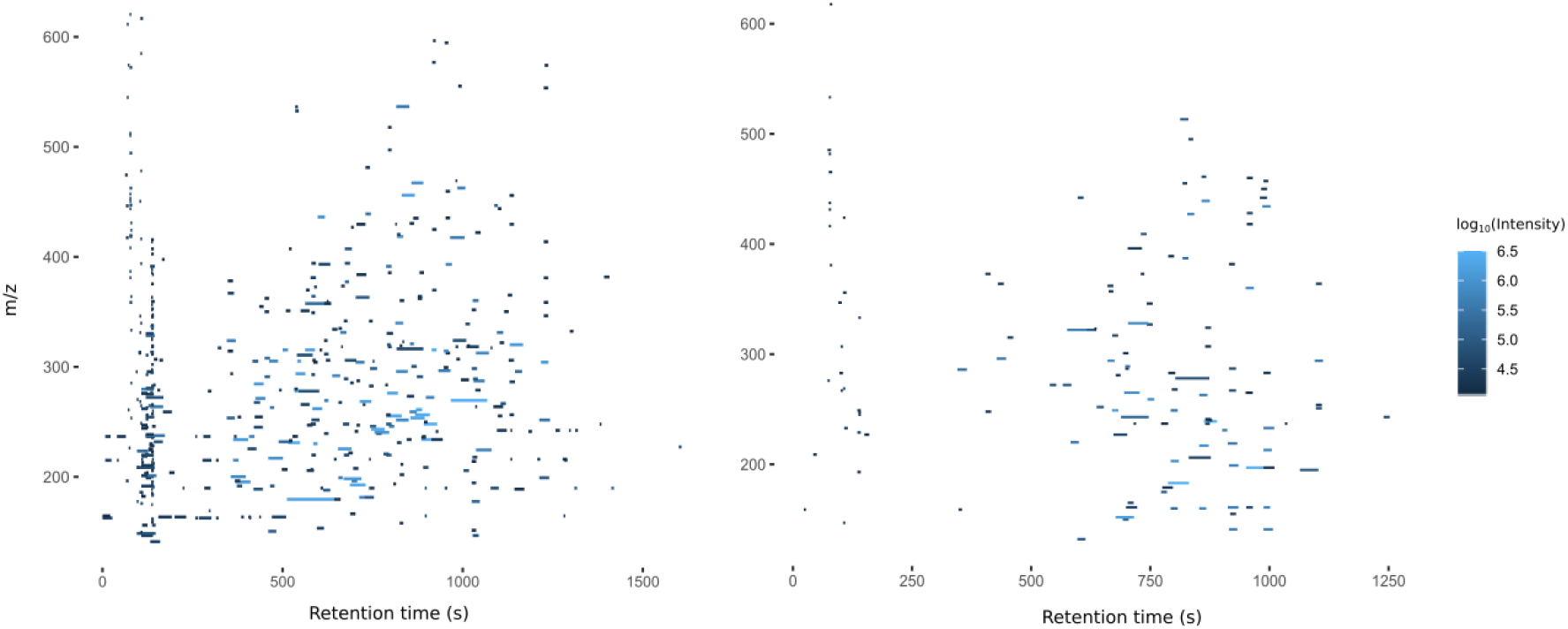
Distribution of inclusion list entries of water in a) positive and b) negative ionization mode after blank subtraction.

**Supplementary Figure 8.**
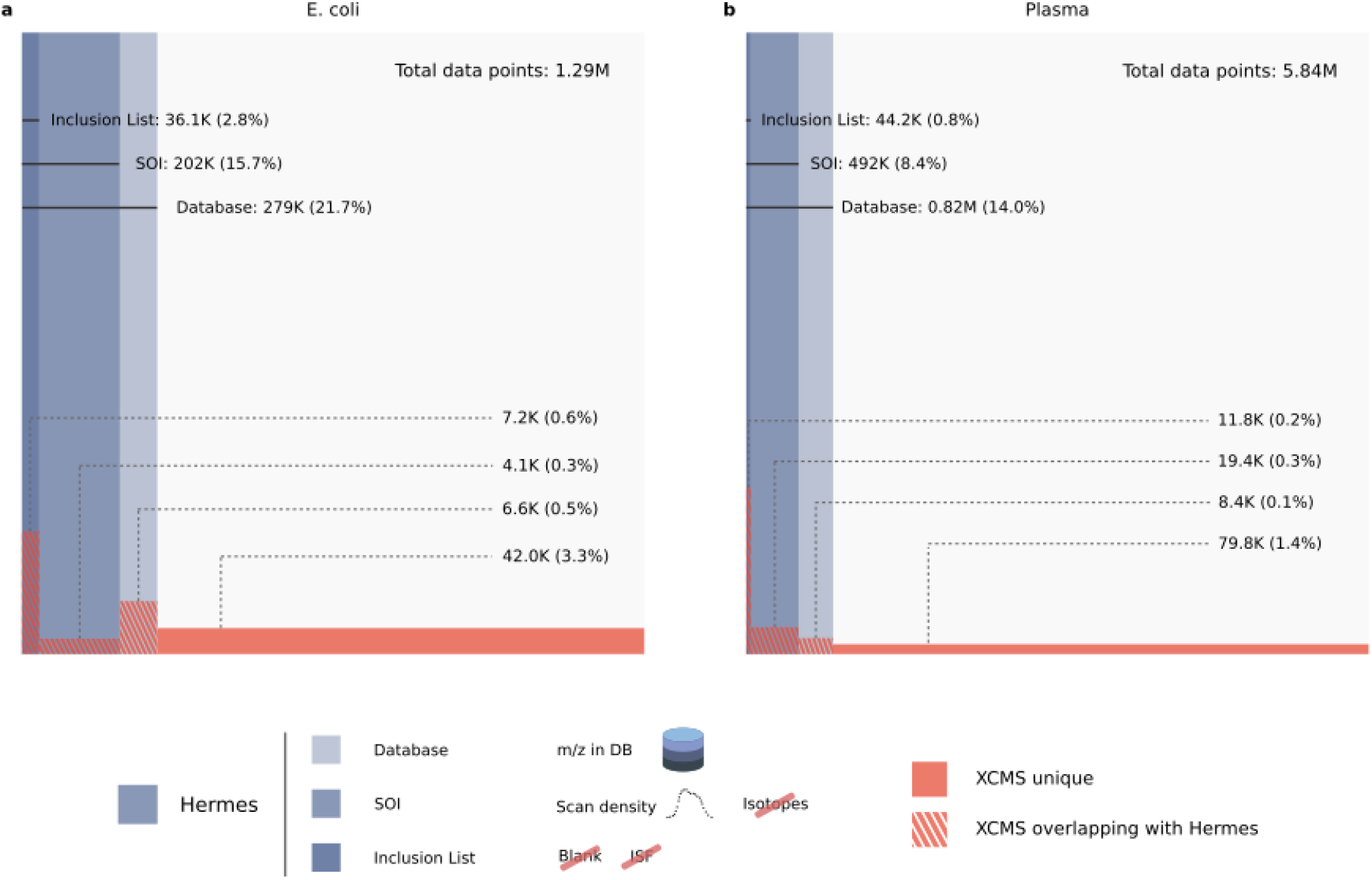
Venn-like diagram of the distribution of negative ionization LC/MS1 data points in different steps of the HERMES workflow and XCMS peak-associated points. a) *E. coli* and b) human plasma extract. Database: Refers to all data points whose m/z matches with any m/z calculated from the ionic formula database (including isotopes). SOI: monoisotopic (M0)-annotated data points that are in Database and are also present in a SOI list that does not include blank subtraction nor any filtering. Inclusion List: data points present in Database and SOI kept through the blank subtraction, isotopic filter and ISF removal steps. Percentages refer to the total number of LC/MS1 data points.

**Supplementary Figure 9.**
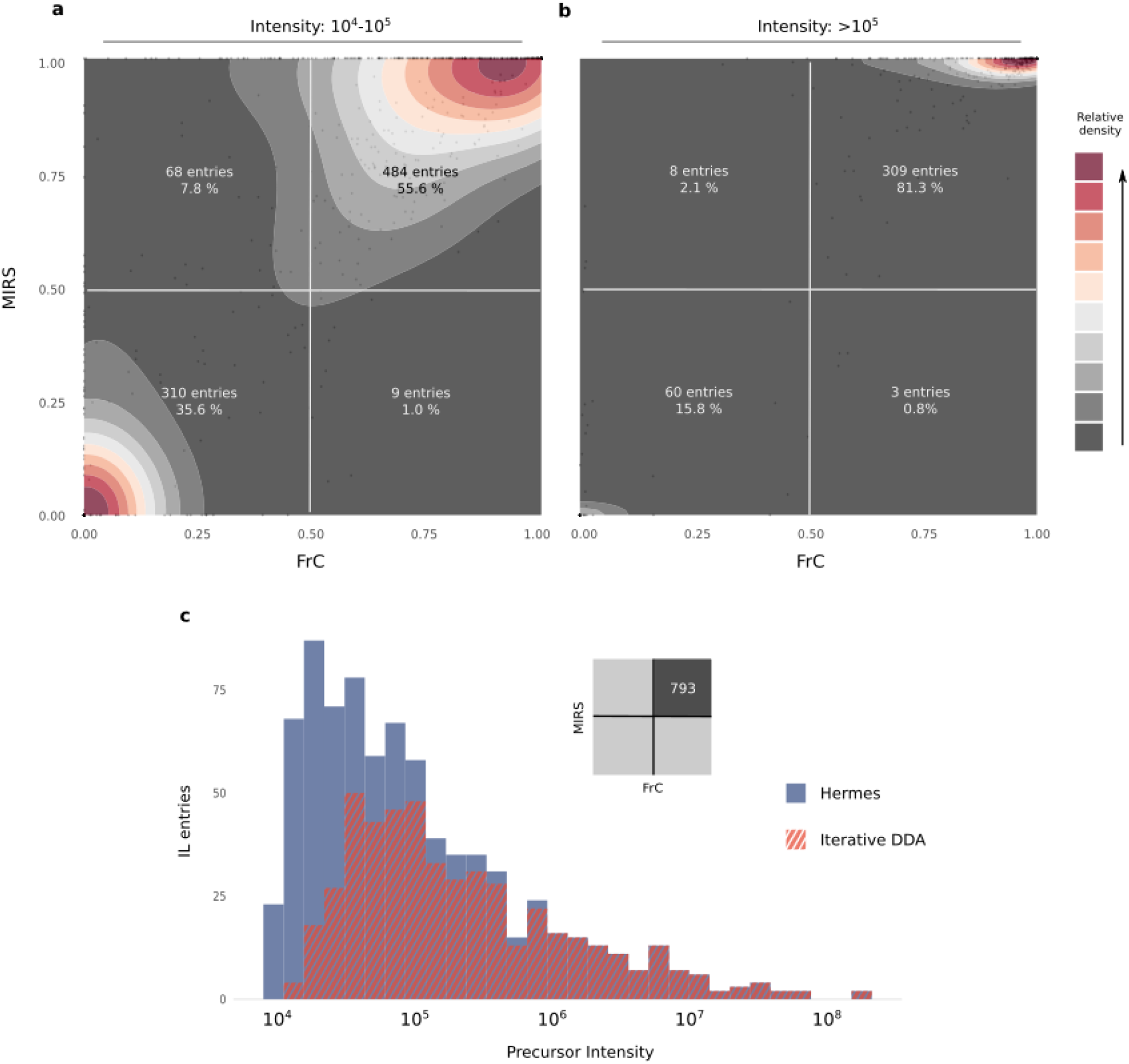
^13^C-enrichment distribution according to the precursor intensity. a) and b) ^13^C-enriched metabolites (FC and MIRS > 0.5) are mainly associated with abundant ions (intensity >10^5^), while unlabeled precursors (FC and MIRS < 0.5) relate more frequently to low abundant ions (intensity between 10^4^-10^5^). c) ^13^C-labeled precursors in iterative DDA corresponded to highly abundant ions that were also covered by HERMES. However, 56% of labeled low abundant ions were not covered by the iterative DDA.

**Supplementary Figure 10.**
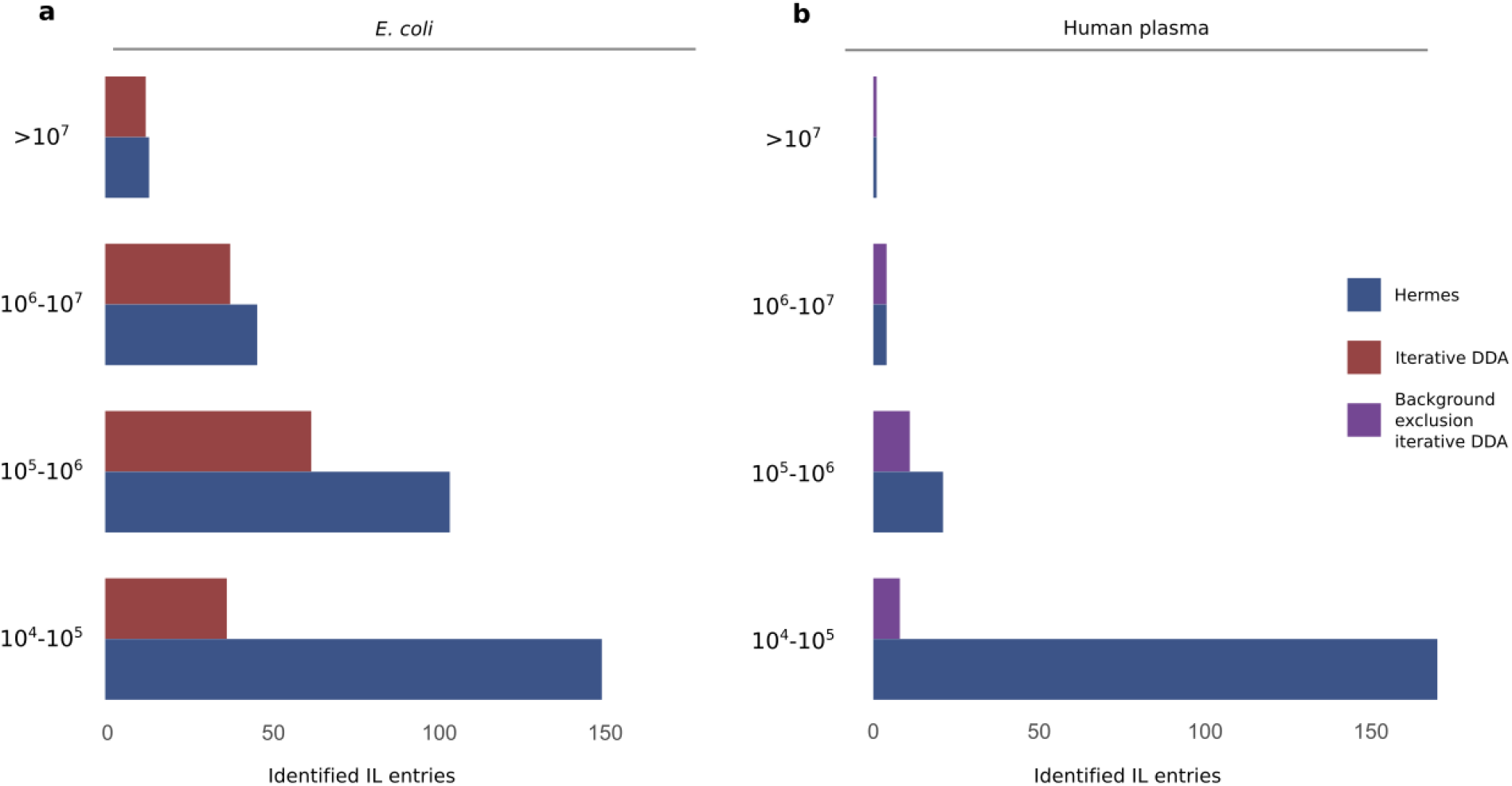
Identified IL entries according to the MS1 precursor intensity. An inclusion list entry is considered identified if at least one MS2 scan associated with it has a compound hit in the reference MS2 database with either cosine score > 0.8 (in-house database from MassBankEU, MoNA, Riken and NIST14 spectra), or Match > 90 and Confidence > 30 (mzCloud). Negative ionization data. a) *E. coli* extract. b) Human plasma extract.

**Supplementary Figure 11.**
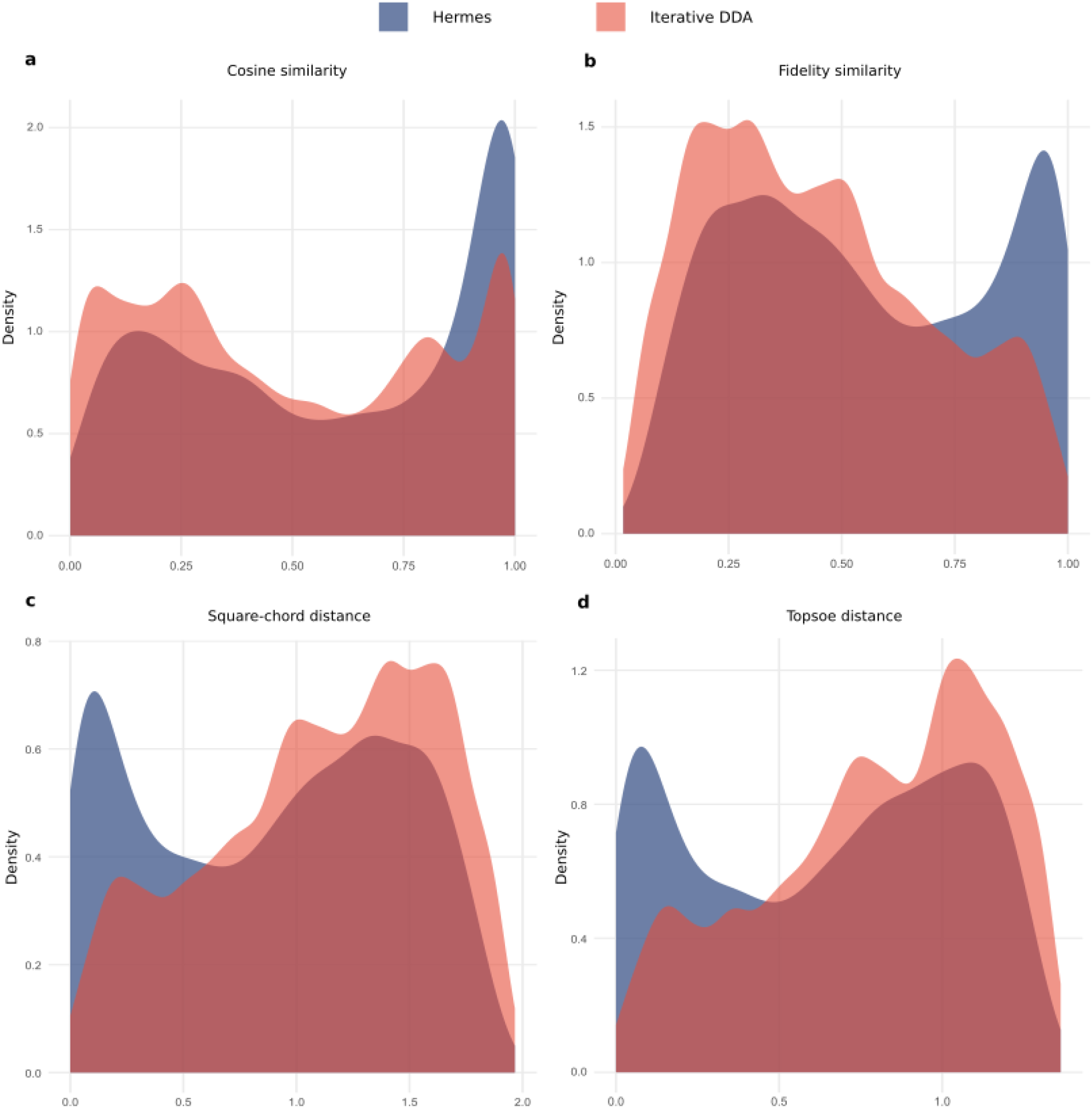
Alternative spectral similarity algorithms and spectrum-spectrum match scores. a) Cosine similarity distribution b) Fidelity similarity distribution. c) Square-chord distance distribution. d) Topsoe distance distribution. A density estimation was calculated with ggplot2 and normalized so that the integral of the curve equals 1. HERMES spectra showed higher similarity scores (a and b) and lower spectral distances (c and d) than DDA spectra.

**Supplementary Figure 12.**
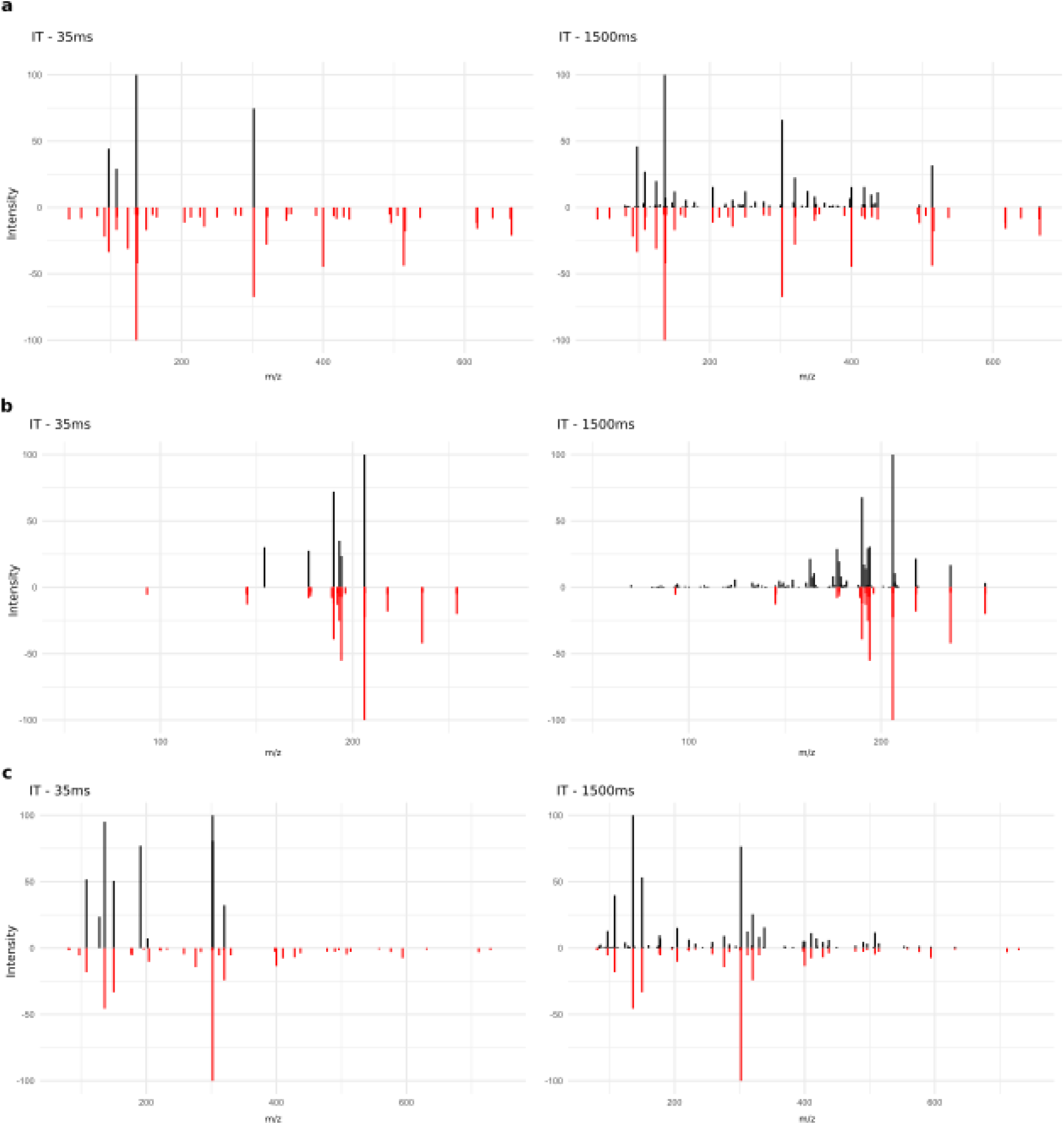
Injection time (IT) comparison (35 ms vs 1,500 ms). Intensity precursor ions <1.0e5. MS/MS (black) of a) NADH, b) Biopterin and c) NADPH against library spectra (red). A higher injection time resulted in richer spectra, with more matching fragments against the reference spectra and overall better matching scores.

**Supplementary Table 1.**
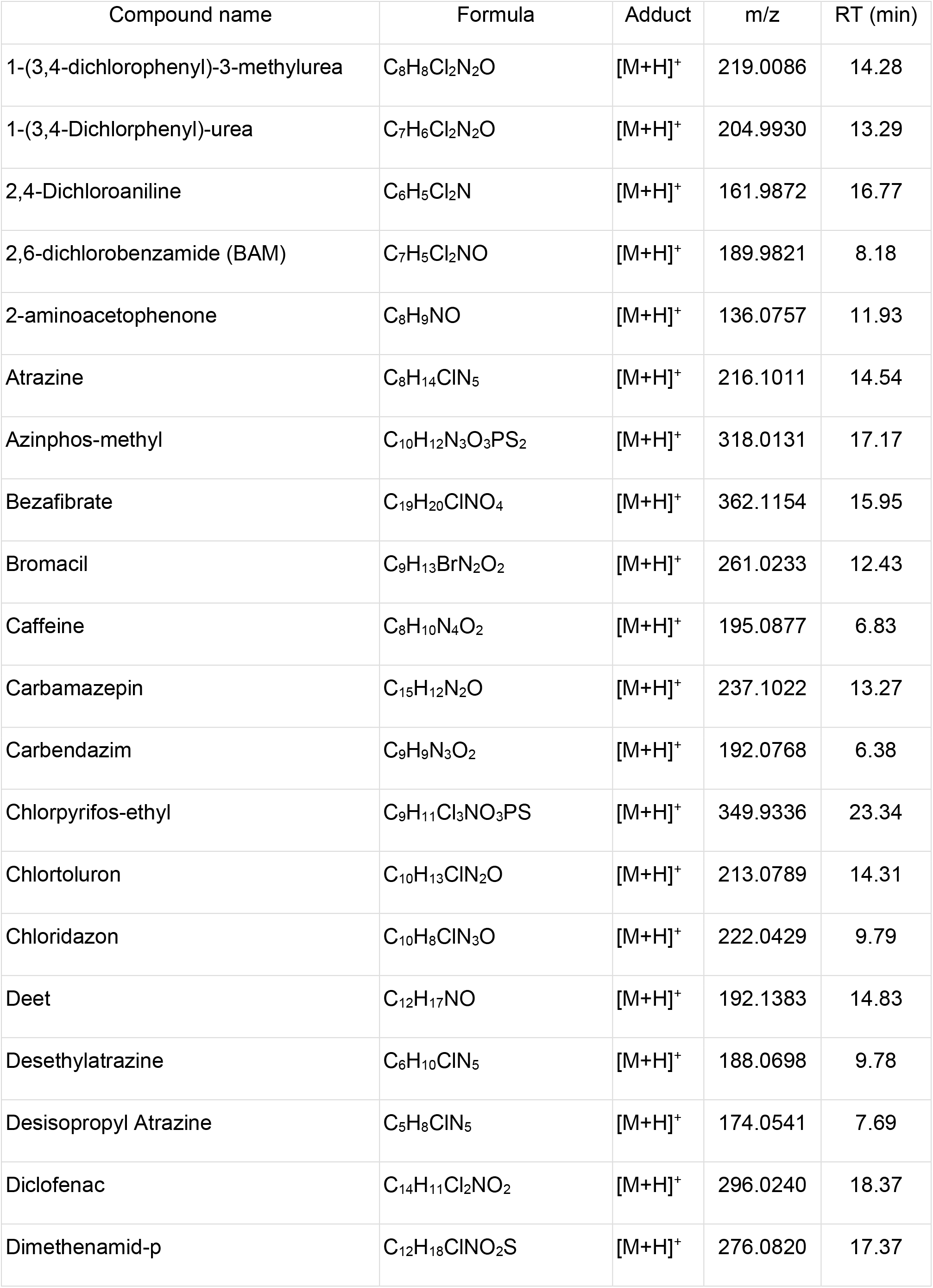
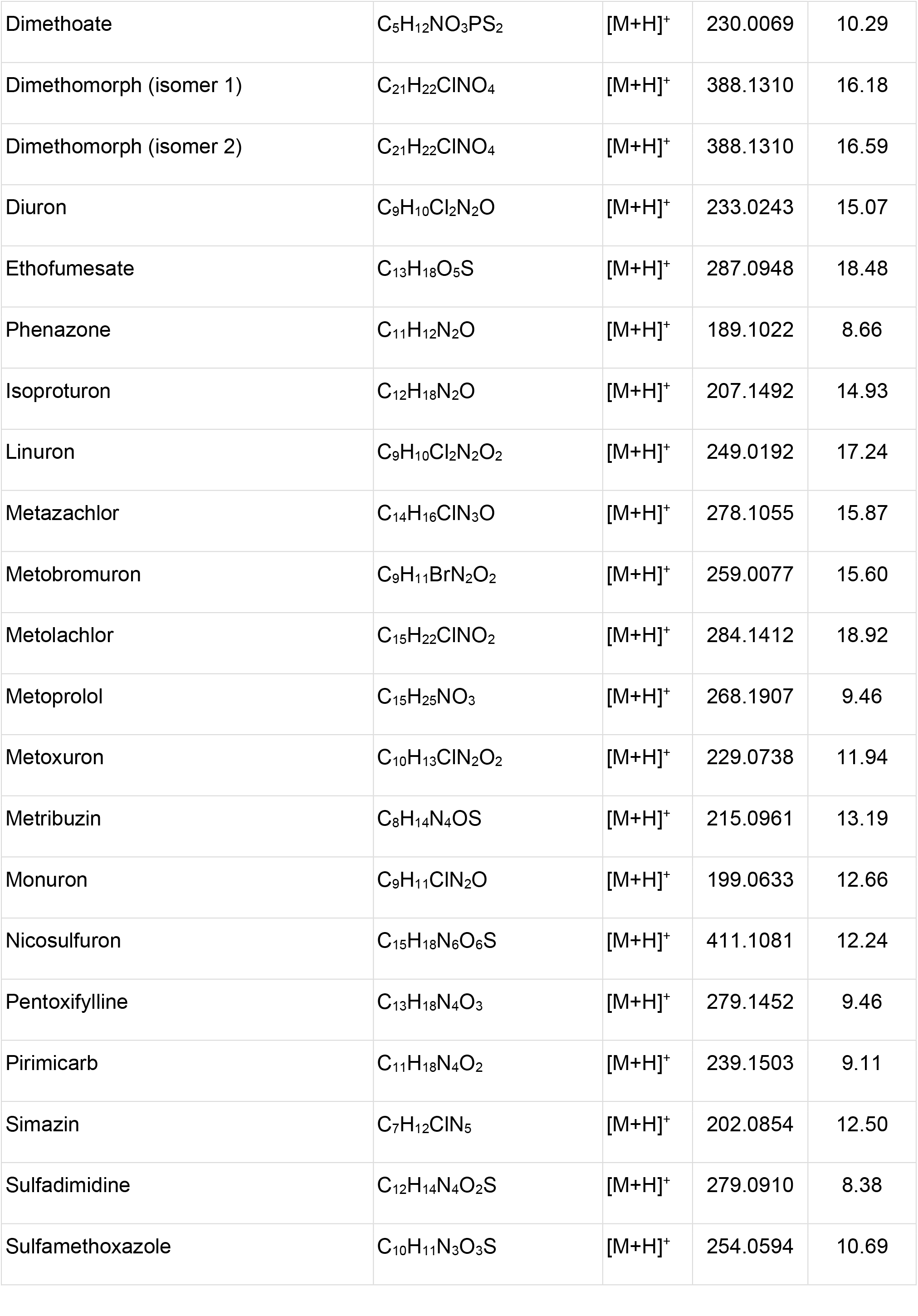
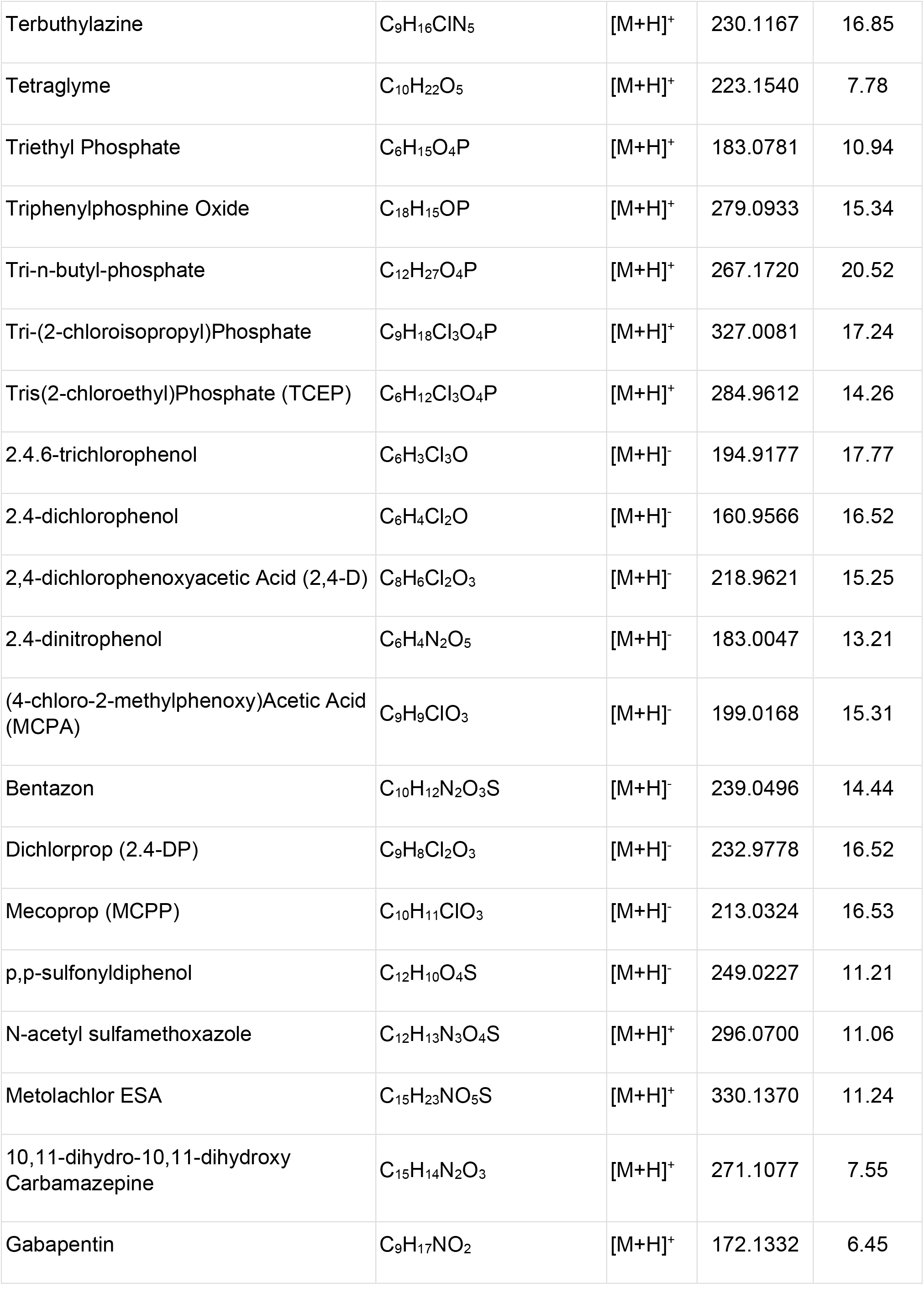
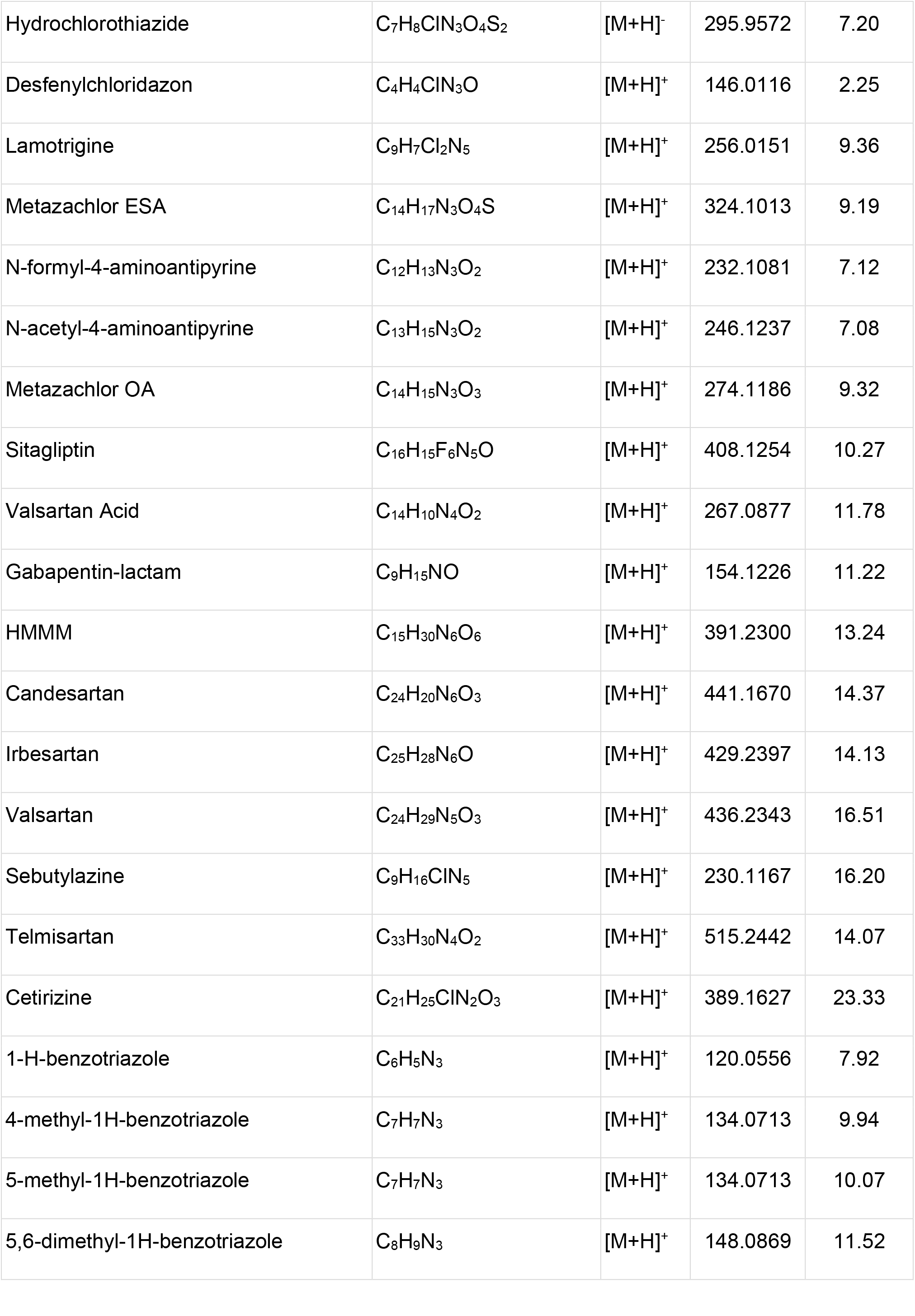
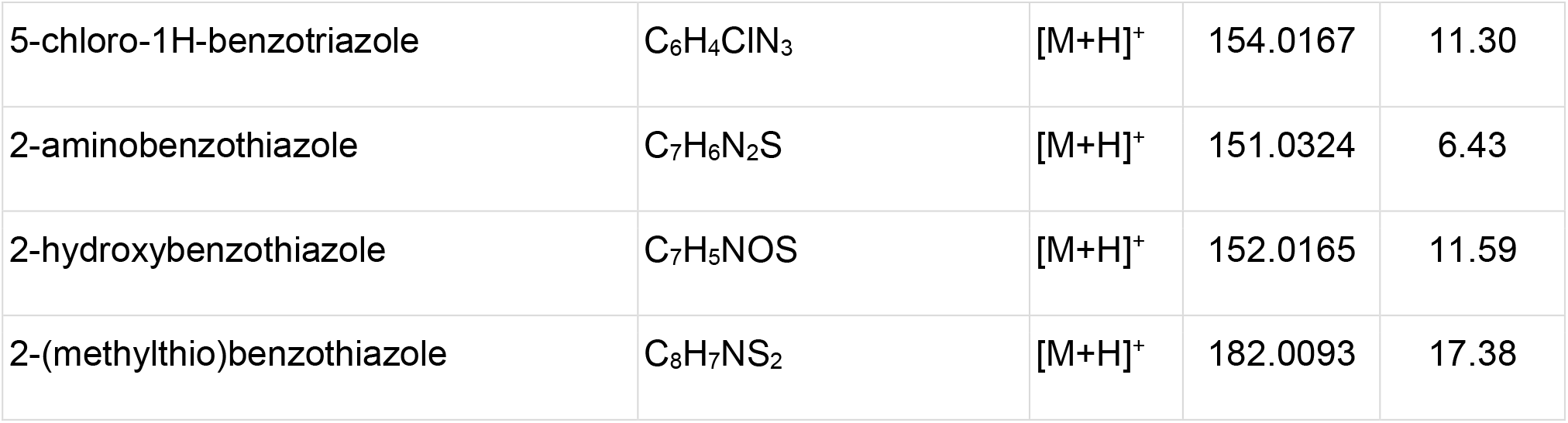
List of spiked compounds

## Notes

https://doi.org/10.5281/zenodo.4581662

### Competing Interest Statement

The authors have declared no competing interest.

